# Hypervoxels: a multidimensional framework for the representation and analysis of neuroimaging data

**DOI:** 10.1101/2022.04.11.485553

**Authors:** Pedro A. Luque Laguna, Ahmad Beyh, Francisco de S. Requejo, Richard Stones, Derek K. Jones, Laura. H. Goldstein, Marco Catani, Steve C.R. Williams, Flavio Dell’Acqua

## Abstract

Most neuroimaging modalities use regular grids of voxels to represent the three-dimensional space occupied by the brain. However, a regular 3D voxel grid does not reflect the anatomical and topological complexity represented by the brain’s white matter connections. In contrast, tractography reconstructions based on diffusion MRI provide a closer characterisation of the white matter pathways followed by the neuronal fibres interconnecting different brain regions. In this work, we introduce hypervoxels as a new methodological framework that combines the spatial encoding capabilities of multidimensional voxels with the anatomical and topological information found in tractography data. We provide a detailed description of the framework and evaluate the benefits of using hypervoxels by carrying out comparative voxel and hypervoxel cluster inference analyses on diffusion MRI data from a neuroimaging study on amyotrophic lateral sclerosis (ALS). Compared to the voxel analyses, the use of hypervoxels improved the detection of effects of interest in the data in terms of statistical significance levels and spatial distribution across white matter regions known to be affected in ALS. In these regions, the hypervoxel results also identified specific white matter pathways that resolve the anatomical ambiguity otherwise observed in the results from the voxel analyses. The observed increase in sensitivity and specificity can be explained by the superior ability of hypervoxel-based methods to represent and disentangle the anatomical overlap of white matter connections. Based on this premise, we expect that the use of hypervoxels should improve the analysis of neuroimaging data when the effects of interest under investigation are expected to be aligned along distinct but potentially overlapping white matter pathways.

## 1. Introduction

Neuroimaging has become an essential tool for the investigation of the living human brain. The analysis of neuroimaging data allows researchers to address multiple questions about the structure and function of the brain and its pathologies. In addition to the individual numerical values of each data point in an image, knowledge of the relative position and topological relationship between different points is fundamental to be able to compare image values across different subjects, and to directly link findings to anatomical regions. This topological information, required for the analysis and interpretation of the image data, depends on the format used for the encoding and representation of the images. In neuroimaging, the most common format consists of multidimensional data matrices representing spatial arrangements of volumetric units called voxels. To fully interpret the information conveyed by a 3D image, each voxel value has to be compared with the values from surrounding voxels (i.e. its topological neighbourhood) and progressively, with the rest of the voxels forming the image. It is the topology of the voxel grid which makes it possible to encode information beyond the numerical values of individual voxels. Without the use of the voxel topology, important information present in the images, such as the underlying anatomy, would be inaccessible and virtually indistinguishable from noise (Fig.1-A).

**[Fig.1].**
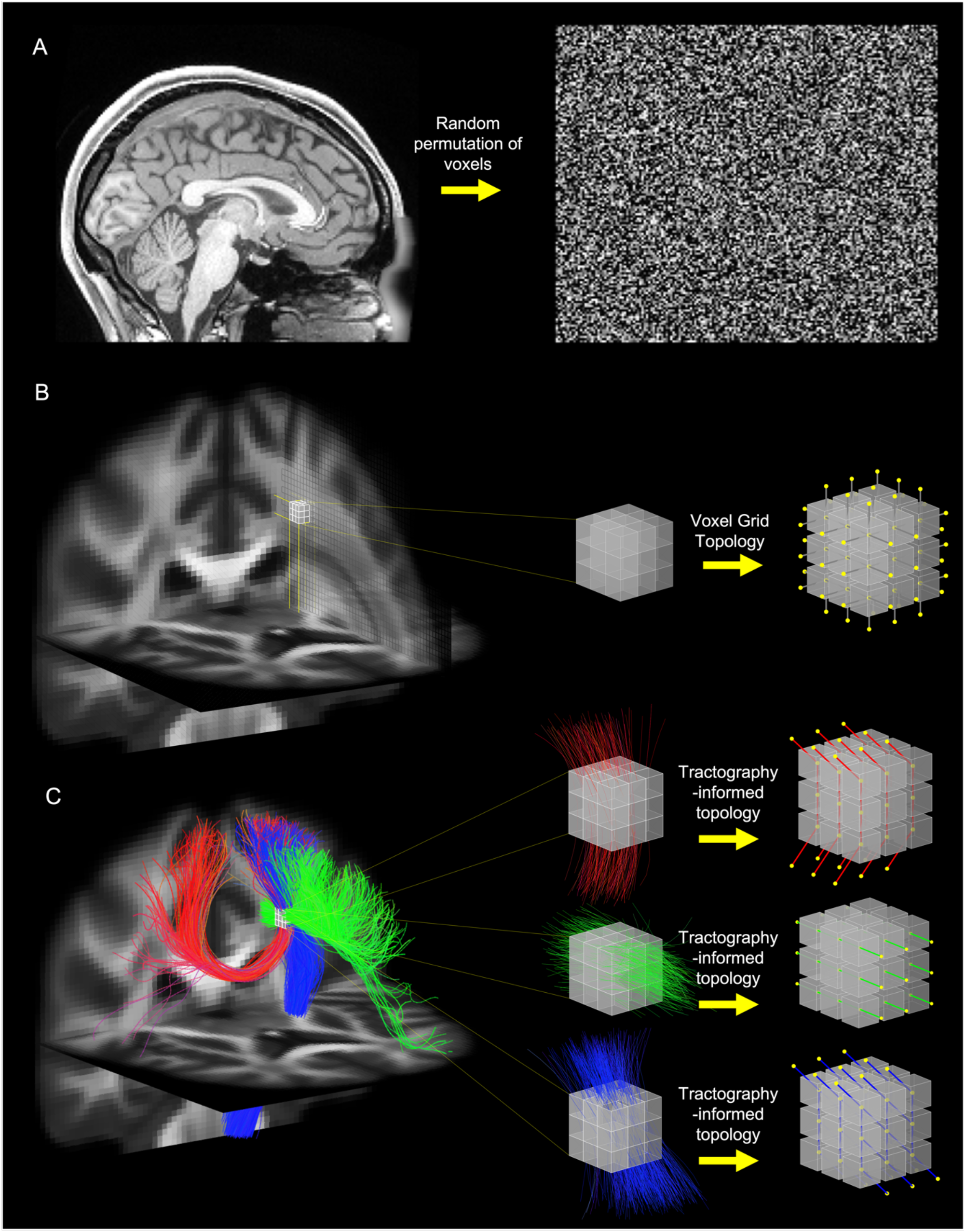
A) Contribution of the voxel topology to the interpretation of an image: The anatomical information is lost after a change in the voxel topology exemplified by a random permutation of the voxel positions (right) showing the critical role played by the topology in the encoding and interpretation of the original image (left). B) Topological properties of a 3D voxel grid: Given an arbitrary neighbourhood of voxels (left) its topology is defined by a pattern of connectivity that remains invariant to rotations and translations within the voxel grid (right). C) Anatomical and topological information provided by tractography: At the macroscale, tractography data provide anatomical pathways used to characterise the structural connectivity of the brain (left). At the voxel level, tractography streamlines can resolve the heterogeneous neuroanatomical complexity represented by axonal fibres and redefine the topological relationship between neighbouring voxels (right).

The topology of a 3D voxel grid corresponds with the structure of a 3D lattice graph that preserves important properties of the 3D Euclidean space. For example, the 3D Euclidean space is isotropic and homogeneous, i.e., its geometrical properties are invariant to rotations and translations. Analogously, the 3D voxel grid shows isotropic connectivity and spatial homogeneity in the sense that the connectivity between neighbouring voxels is defined by a pattern that remains invariant to rotations and translations within the grid (Fig.1-B). This facilitates in great manner the spatial manipulation of 3D data with efficient mathematical computations, which makes the voxel grid a convenient framework for the representation of 3D images using a format that is also independent of the object depicted. The use of the voxel grid by early computational methods for the anatomical registration and the statistical analysis of brain images (Alpert et al. 1990; K. J. Friston, Jezzard, and Turner 1994; K. J. Friston et al. 1995; Worsley et al. 1992; Wright et al. 1995) paved the way for the development of publicly available software packages for neuroimaging analysis based on voxels, such as, FSL (Jenkinson et al. 2012), SPM (Ashburner 2012), and AFNI (Cox 2012). However, it is often the case that the information of interest present in the images involves topological aspects of the brain anatomy that do not align with the structure of the 3D voxel grid, in which case its use may not be the most appropriate for an effective analysis of the imaging data.

For example, to increase their sensitivity, many methods use spatial inference techniques that identify “blobs” or “clusters of voxels” associated with the presence of an effect of interest in the data over an extended region of space. They provide supporting evidence of the implication of the underlying brain regions in the investigated phenomena. In most cases, however, it is not possible to ensure the anatomical specificity of the results because the interconnected voxels that make each cluster can span multiple brain regions or/and anatomical structures. Also, individual voxels can represent more than one brain structure, sometimes due to the coarse resolution of the images, but often as a consequence of the intrinsic anatomical overlap between brain structures that the voxel topology cannot resolve. The anatomical uncertainty of the voxels is even amplified in methods like TBSS that project the voxel values into an image skeleton to boost the sensitivity of the analysis (Bach et al. 2014; S. M. Smith et al. 2006). In all these scenarios, the mismatch between the voxel topology and the brain anatomy could be contributing to the apparent trade-off between the sensitivity of the analyses and the anatomical specificity of their results.

The other topological aspect of the brain anatomy that differs from the voxel grid is its structural connectivity. In contrast to the isotropy and homogeneity that characterise the connectivity of the voxel grid, the alignment of the neural connections across the brain is neither isotropic nor homogeneous. For example, in grey matter regions, the orientation of the neural fibres reflects the organisation of neurons in layers and columns. Relative to the cortex, connections are vertical between neurons in the same column, and mostly horizontal between neurons in the same layer (Schnepel et al. 2015). When we look at the white matter, we find myelinated axons arranged into bundles that overlap or intersect each other forming complex patterns of parallel or crossing fibres that also change across different regions [Castro et al. (2005); Silva and Andrade (2016)]. In addition, the intricacy and spatial overlap of the neural connections reflect a topological complexity in the brain that exceeds anything achievable by the connectivity of a 3D voxel grid [Bassett et al. (2010); Pineda-Pardo et al. (2015)], however useful such information would be to the analysis of the images.

In this context, diffusion MRI tractography emerges as a neuroimaging technique that aims, among other things, to characterise the structural connectivity of the living human brain. By analysing the patterns of water diffusion across brain voxels, this technique generates streamlines that aim to represent the white-matter pathways of neuronal axons connecting different brain regions. This ability to display white-matter connections has found several clinical applications (Farquharson et al. 2013; Yamada et al. 2009; Nimsky, Ganslandt, and Fahlbusch 2006) and has advanced the study of the brain’s anatomy and its structural connectivity (Catani et al. 2002; Johansen-Berg and Rushworth 2009). Tractography has also made an important contribution to the analysis of neuroimaging data with its application to tractometry (Bell et al. 2011; Yeatman et al. 2012) and to connectomic analyses (Hagmann 2005; Hagmann et al. 2010; Sporns, Tononi, and Kötter 2005). In tractometry, the streamlines are used to identify white-matter pathways in voxel-based images from which quantitative values are extracted, averaged and analysed. This can be done for specific tracts in the fashion of region of interest (ROI) analyses to increase the statistical power and the anatomical specificity of the results (Jones and Nilsson 2015), or across the entire white matter using analysis methods based on automated tractography clustering techniques (Siless et al. 2020; Zhang, Wu, Ning, et al. 2018). In a connectomic analysis, the brain’s structural connectivity is modelled as networks of grey matter regions (nodes) interconnected by the streamlines (edges) that are analysed as graph-theory objects in terms of connectivity strength or other topological properties (Fornito, Zalesky, and Breakspear 2015). In all these analyses, incorporating the topological information provided by the streamlines entails exchanging the spatial resolution of the voxels with the “anatomical” resolution associated with the white-matter tracts or with the interconnected grey matter regions. However, the streamlines also provide a geometrical approximation to the architectural complexity associated with the local fibres (F. Dell’Acqua and Tournier 2019; Jeurissen et al. 2019), and their connectivity applies to all the voxels intersected by the streamlines and not just to the grey matter regions interconnected by them. This information can be used to increase the anatomical specificity provided by the voxels and to redefine the topological relationship between them (Fig.1-C).

*In this paper, we introduce hypervoxels as a new framework for the representation and analysis of neuroimaging data that combines the spatial encoding capabilities of multidimensional voxels with the geometrical and topological properties of tractography streamlines*. We extend the 3D voxel grid with additional spatial dimensions that encode *local* and *global* information about the geometry of the streamlines that intersect each voxel. The newly defined hypervoxel grid allows us to represent the streamline space in a voxel-compatible framework that maintains the spatial resolution of the images. At the same time, we go beyond the traditional connectivity of the regular voxel grid by providing a new anatomically inspired topology directly based on the geometry of fibre bundles. This new topology identifies hypervoxels in the hypervoxel grid that are either connected or adjacent to each other according to the trajectories of the streamlines from a given tractography dataset. The result is a hypervoxel encoding, i.e., a high-dimensional voxel-like topological discretization of the “anatomical” space represented by a collection of tractography streamlines. We provide examples on how to perform this hypervoxel encoding and how to use it in the analysis of neuroimaging data. For that, we first construct a hypervoxel template using tractography data obtained from a group of healthy individuals. Then, we provide the hypervoxel implementation of two standard methods used in the statistical analysis of neuroimaging data: cluster-level inference and threshold-free cluster enhancement (TFCE). Finally, we use these two methods to analyse MRI data acquired on previous neuroimaging studies and compare the results obtained using either the voxel or the hypervoxel version of each method.

## 2. Methods

### 2.1. Hypervoxels: local information

Hypervoxels extend the space represented by a voxel grid with extra dimensions that represent additional geometrical information about the streamlines. We start by considering the Position-Orientation (*P-O*) space (Tuch 2002; Hagmann 2005) representing all possible combinations of 3D positions *and* local angular orientations.

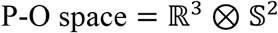

In P-O space, every streamline ***s*** has associated a sequence 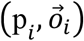 of vertex positions and local angular orientations where each local orientation 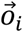 corresponds to the unit vector tangent to the trajectory of **s** at each vertex position 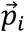 (Fig.2 - centre-up).

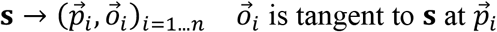

To encode this combined information of positions and orientations, we extend a 3D voxel position grid **[X, Y, Z]** with an additional dimension ***O*** that represents an orientation in a particular discretisation of the sphere (Fig.2 - centre-down). The result is a tessellation of P-O space in the form of a hypervoxel grid **[X, Y, Z, O]** that allows us to represent each streamline as a sequence of hypervoxels defined by their corresponding hypervoxel coordinates [*x*_*i*_, *y*_*i*_, *z*_*i*_, *o*_*i*_].

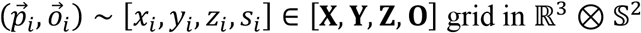

The P-O hypervoxel with coordinates [*x*_*i*_, *y*_*i*_, *z*_*i*_, *o*_*i*_] locally represents all streamlines intersecting voxel [*x*_*i*_, *y*_*i*_, *z*_*i*_] at an angle within the spherical section given by the grid element [*o*_*i*_]. This constitutes a “one-to-many” correspondence between each hypervoxel and the locally represented streamlines. Here, the granularity of the angular encoding is determined by the resolution of the hypervoxel grid in the **[O]** dimension. For example, a homogeneous parcellation of the sphere in 100 regions would yield an angular resolution of 4*π*/100 steradians per voxel. This is the analogy of the granularity of the spatial encoding given by the voxel resolution (i.e. 1,2,3, etc mm).

**[Fig.2].**
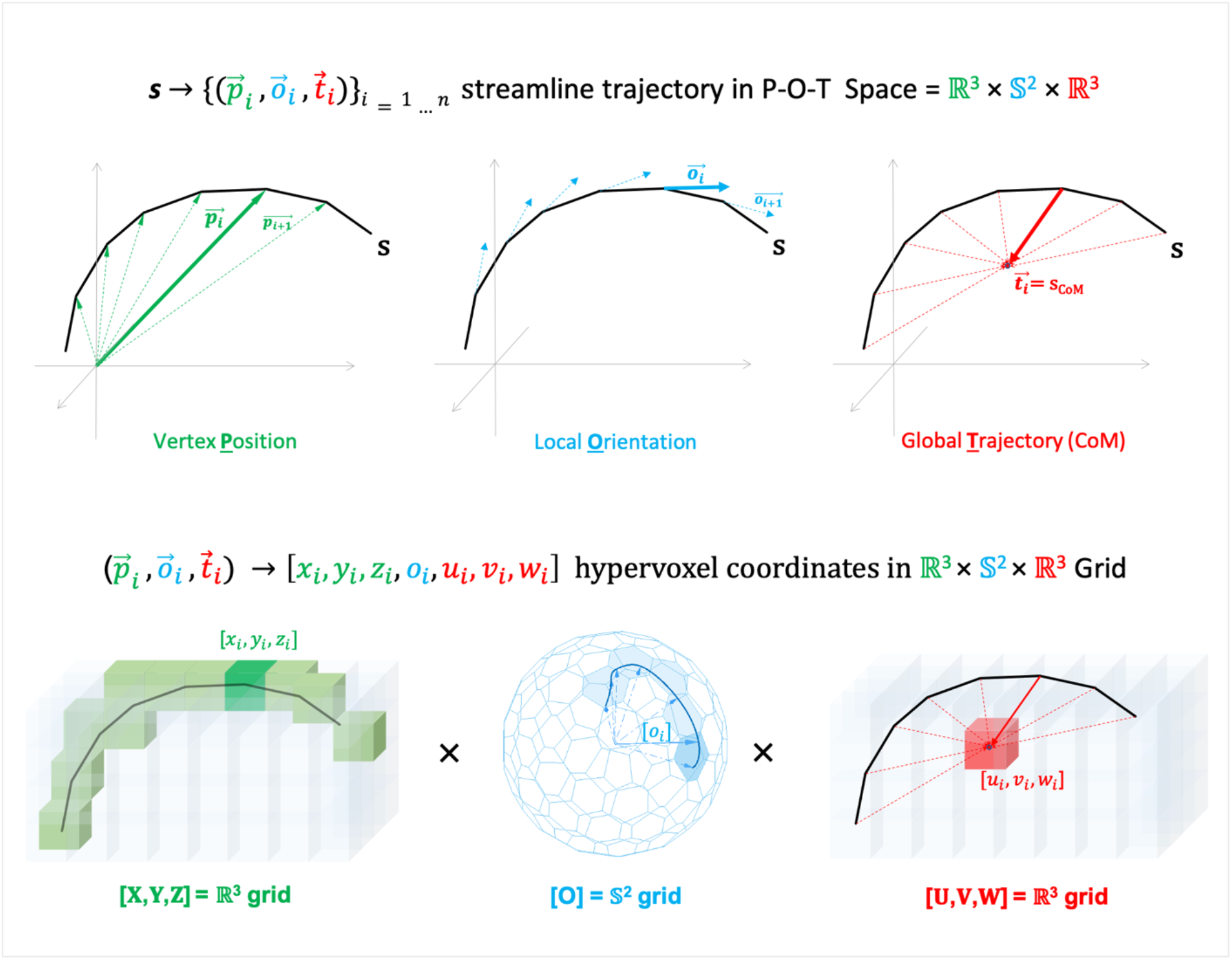
Each streamline corresponds with a trajectory of points in P-O-T Space given by vertex positions, local orientations and a global trajectory parameter defined by each streamline’s centre of mass. The hypervoxel grid provides a discretisation of P-O-T Space so each combination of position, orientation and trajectory parameter corresponds with an unique hypervoxel in the hypervoxel grid.

### 2.2. Hypervoxels: global information

Together with local information we are also interested in using hypervoxels to capture information about the trajectories followed by the streamlines outside the voxel. Such information can be encoded as a global parameter using additional hypervoxel dimensions. One example is to encode the centre of mass (CoM) of each streamline as a global trajectory parameter 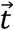 that summarises the entire trajectory of each streamline. The motivation for choosing this parameter is that streamlines connecting the same pair of brain regions tend to follow similar trajectories throughout the brain and therefore have similar CoM. The inclusion of the parameter 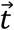 in the definition of each hypervoxel allows us to locally represent streamlines that follow different trajectories outside the voxel as different hypervoxels. Now, each streamline ***s*** corresponds with a sequence of vertex positions, local orientations, and global trajectory parameters in a Position-Orientation-Trajectory (POT) space.

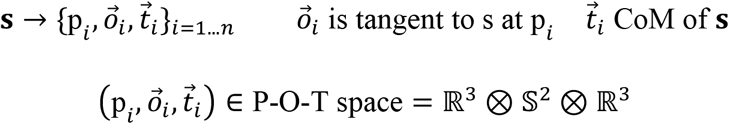

To represent P-O-T space, we extend again the hypervoxel grid with additional dimensions **U, V** and **W** to encode the global trajectory parameter 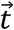 in ℝ^3^. In the extended hypervoxel grid **[X, Y, Z, O, U, V, W]** each streamline is represented by a sequence of 7D hypervoxels.

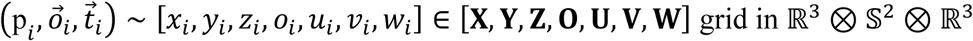

Each hypervoxel now distinguishes between streamlines that intersect a given voxel with distinct local orientation and follow different global trajectory (measured by the CoM) outside the voxel. As for the local orientation, the granularity of the encoding of the global trajectory will be determined by the resolution chosen for the hypervoxel grid.

### 2.3. Alternative global encodings

The combination of local orientation and global trajectory parameters allows the hypervoxel grid to disentangle the anatomical complexity represented by the streamlines intersecting each voxel. Nevertheless, other streamline information could also be used for the definition of the hypervoxels. For example, we could encode the position of the streamline’s termination points as they provide specific information regarding the end-to-end connectivity in the brain.

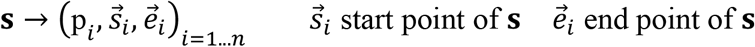

In this case, the space and the corresponding hypervoxel grid would have 10 dimensions to encode vertex position (**[X, Y, Z]**), local orientation (**[O]**), start point (**[SX, SY, SZ]**) and end point (**[EX, EY, EZ]**) of each streamline.

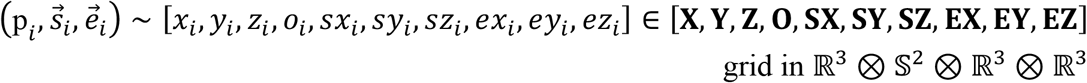

This alternative hypervoxel encoding would allow the differentiation at the voxel level between intersecting streamlines that interconnect different brain regions, information that can be particularly useful for connectomics applications. In general, multiple and distinct global (and local) parameters can be included in the hypervoxel framework.

### 2.4. Hypervoxel encoding of tractography data

Given a tractography dataset and a hypervoxel grid defined by rules to encode some hypervoxel parameters (e.g. local orientation, CoM, etc), the associated tractography hypervoxel encoding is the smallest set of hypervoxels from the hypervoxel grid that represents all the streamlines in the tractography dataset. The following algorithm describes the hypervoxel encoding of a given tractography dataset *{s}* using the local orientation and the CoM as hypervoxel parameters:

CREATE_HYPERVOXEL_ENCODING

- Inputs: TractographyDataset, HypervoxelGrid, Output : HypervoxelEncoding
- For each streamline **s** in TractographyDataset:
  – For every vertex **v** in **s**:
    - Get the voxel coordinates [x,y,z] corresponding to the position of **v**.
    - Calculate the local angular orientation 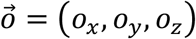 defined by the unit vector tangent to **s** at vertex **v**.
    - Obtain the local orientation coordinate [o] corresponding to 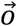 in the discrete parcellation **[O]** of the sphere.
    - Calculate the global trajectory parameter 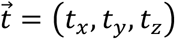 at **v** defined by the relative position of the CoM of the streamline **s** respect to the vertex **v**.
    - Obtain the global trajectory coordinates [u,v,w] by discretising the global trajectory parameter 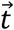 along the **[U, V, W]** dimensions of the HypervoxelGrid.
    - Add the hypervoxel defined by coordinates **hv**=[x, y, z, o, u, v, w] to the sequence of hypervoxels {*hv(s)*} representing the streamline **s**.
  – Add the hypervoxels in {*hv(s)*} to the HypervoxelEncoding.
- Remove any duplicated hypervoxels from the HypervoxelEncoding

The encoded hypervoxels represent only a fraction of the total number of potential hypervoxels in a hypervoxel grid, even for a whole-brain tractography dataset. Also, the encoded hypervoxels are more sparsely distributed (in hypervoxel space) than the voxels intersected by tractography streamlines (in 3D space). The sparsity of the hypervoxel encoding reflects the fact that the streamlines intersecting each voxel represent only a small fraction of the potential trajectories available within the hypervoxel grid.

### 2.5. Hypervoxel topology

Once the hypervoxel encoding is applied to a set of streamlines, we can define a new topology on the encoded hypervoxels that better reflects the connectivity and geometry associated with the streamlines. For that purpose, we take inspiration from the geometry of *mathematical* fibre bundles (Ivancevic and Ivancevic 2007) that resemble the *anatomical* fibre bundles associated with white matter tracts. A mathematical fibre bundle corresponds *locally* with the product of two spaces referred to as the base manifold and the fibre. The local geometry at each point is given by the direct sum of two vectorial components, one vertical and one horizontal (Kolář, Slovák, and Michor 1993). The vertical component is the vertical bundle formed by the vectors that are tangent to the fibres. Its complement is the horizontal bundle given by an Ehresmann connection (Ehresmann 1950). The Ehresmann connection is a mathematical object that allows the definition of horizontal sections in the fibre bundle and the parallel transport of vectors across the horizontal sections and along the fibres (Fig.3).

**[Fig.3].**
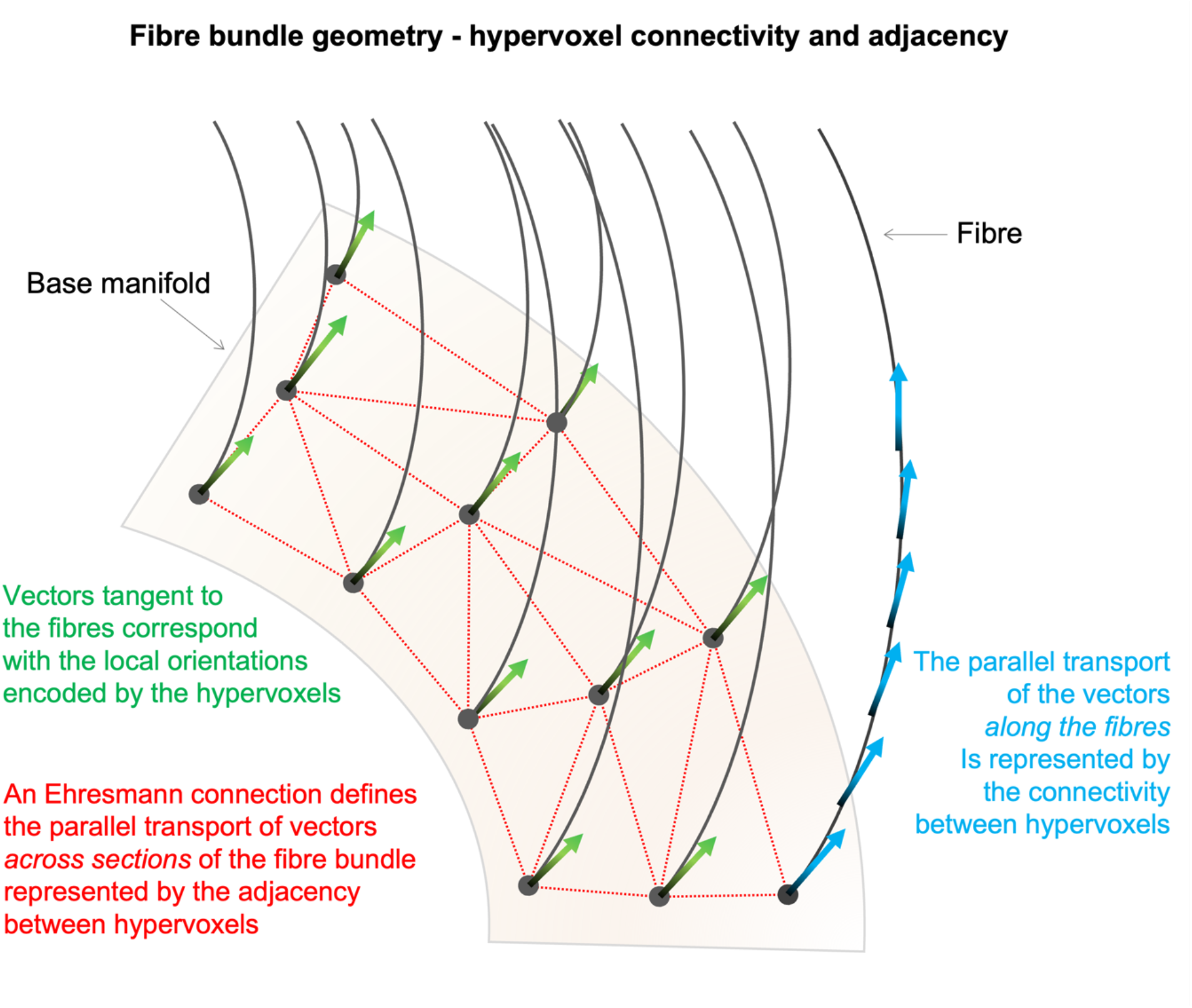
Fibre bundle geometry. The connectivity and adjacency between hypervoxels is inspired by the geometry of fibre bundles. The hypervoxel connectivity reflects the vectorial parallel transport along the fibres. The adjacency between hypervoxels is inspired by the parallel transport of vectors across horizontal sections of the bundle, as defined by an Ehresmann connection.

When we identify a bundle of streamlines with a mathematical fibre bundle, the local orientations of the streamlines encoded by the hypervoxels correspond with the tangent vectors that form the vertical bundle, while the cross sections of parallel axonal fibres corresponds with the horizontal sections defined by an Ehesmann connection. Based on this analogy, we use the parallel transport of vectors along the fibres and across horizontal sections of a fibre bundle to motivate the following complimentary criteria of connectivity and adjacency between hypervoxels:

#### Hypervoxel connectivity

two hypervoxels are longitudinally connected if there is a streamline that is locally encoded by both hypervoxels (i.e. they are “intersected” by the same streamline) and they are not separated from each other more than a predefined distance threshold used to account for tractography propagation errors. This longitudinal connectivity criterion gives rise to a hypervoxel along-tract connectivity matrix *HV*_*c*_, a square binary matrix indicating which hypervoxels are interconnected to one another by at least one streamline and separated along the streamline no more than the specified distance threshold.

#### Hypervoxel adjacency

two hypervoxels are radially adjacent if their “voxel” coordinates are adjacent to each other and they have the same coordinates for the rest of hypervoxel parameters (Fig.4). The corresponding hypervoxel radial adjacency matrix *HV*_*a*_ indicates which hypervoxels are radially adjacent to each other in the hypervoxel encoding.

**[Fig.4].**
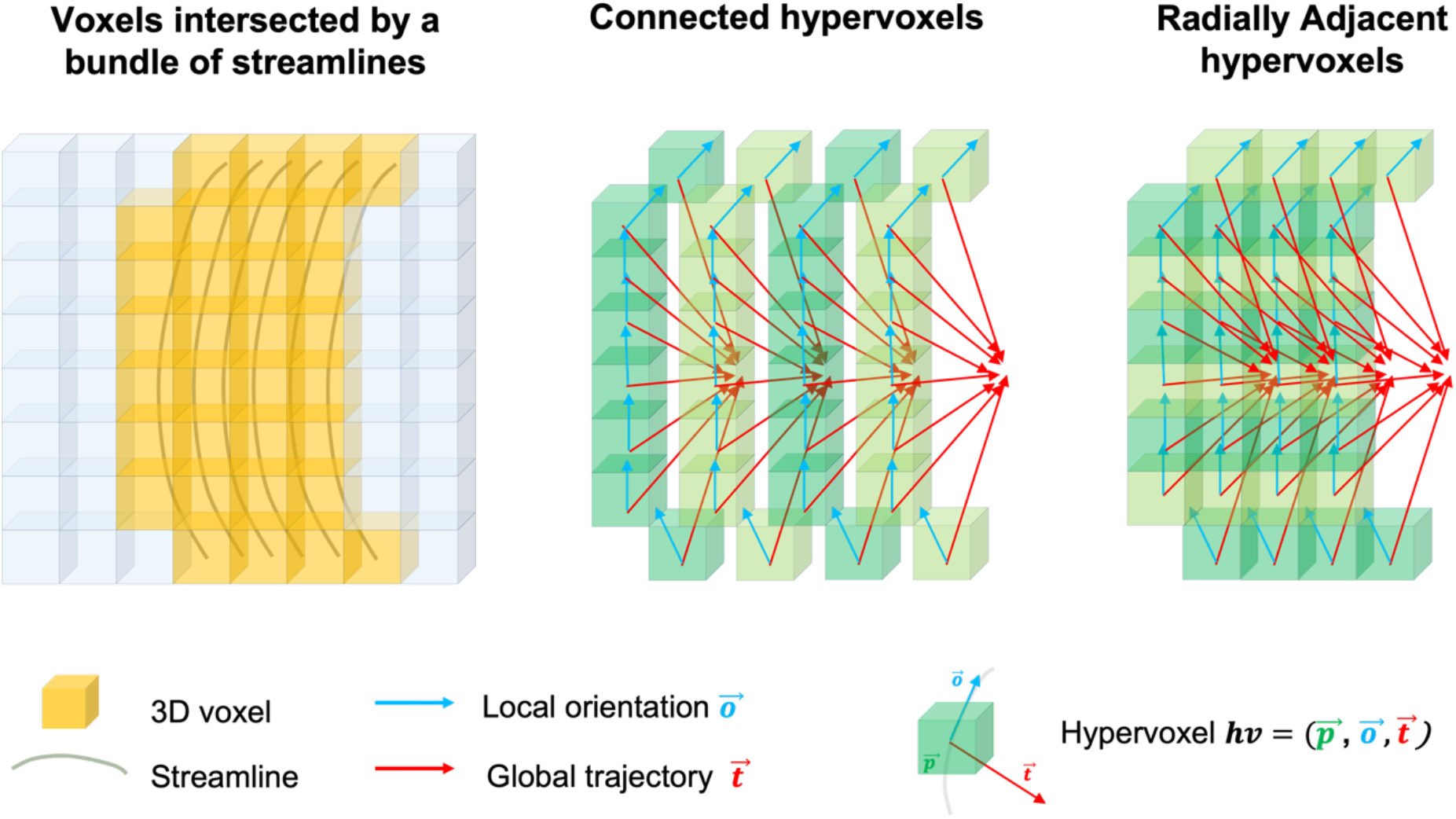
Hypervoxel topology. The intersection of voxels by a bundle of parallel streamlines produces a collection of hypervoxels topologically related with each other. Connected hypervoxels: those hypervoxels representing voxels intersected by streamlines that follow the same trajectory within the bundle. Radially adjacent hypervoxels: hypervoxels representing adjacent voxels intersected by streamlines following parallel trajectories within the bundle.

The hypervoxel connectivity and adjacency matrices provide the hypervoxel encoding with a new topological structure that reflects the organisation of the streamlines as fibre bundles. Here, connected hypervoxels form the vertical component of a bundle (i.e. the fibres), while adjacent hypervoxels form the horizontal component of the bundle (i.e. the cross-sections). This new topology constitutes a departure from the connectivity of the voxel grid where each voxel is “isotropically” connected to all the surrounding voxels (Fig.1-B). Following a hypervoxel encoding, most hypervoxels will be “disconnected” from surrounding hypervoxels in the grid except from those that are part of the same fibre bundle, either as connected or as adjacent hypervoxels. The basic principles of the hypervoxel topology are schematically illustrated in (Fig.4). For a more comprehensive visualisation of the hypervoxel topology following the encoding of a whole-brain tractography dataset, see (Fig.7)-B-D-E.

**[Fig.5].**
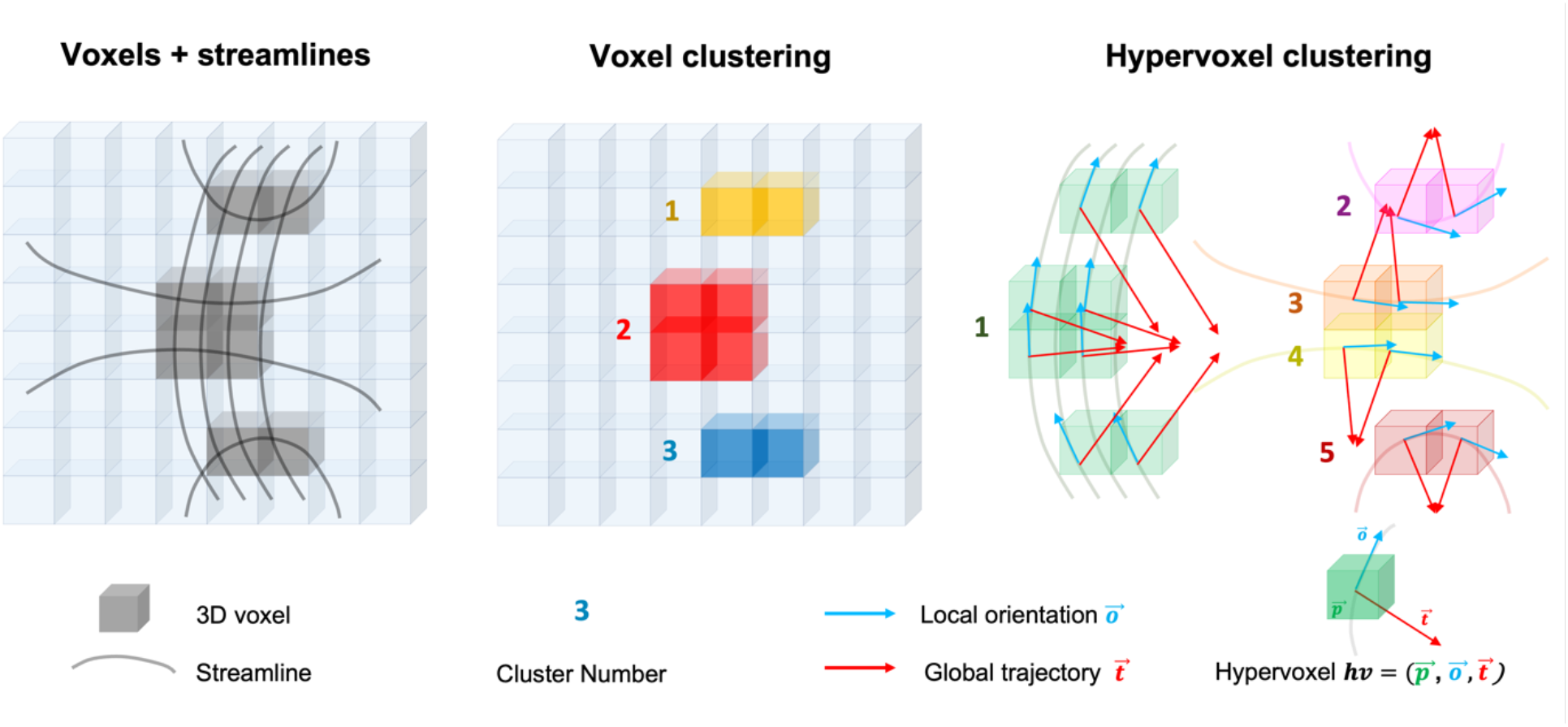
Clustering of voxels vs clustering of hypervoxels. Voxel clustering produces 3 clusters defined exclusively by the spatial adjacency between voxels. Hypervoxels allow the disentangling of regions characterised by the presence of crossing and kissing fibres into separate clusters, each one representing a different group of interconnected or adjacent hypervoxels that reflect the fibre bundle topology associated to the streamlines.

**[Fig.6].**
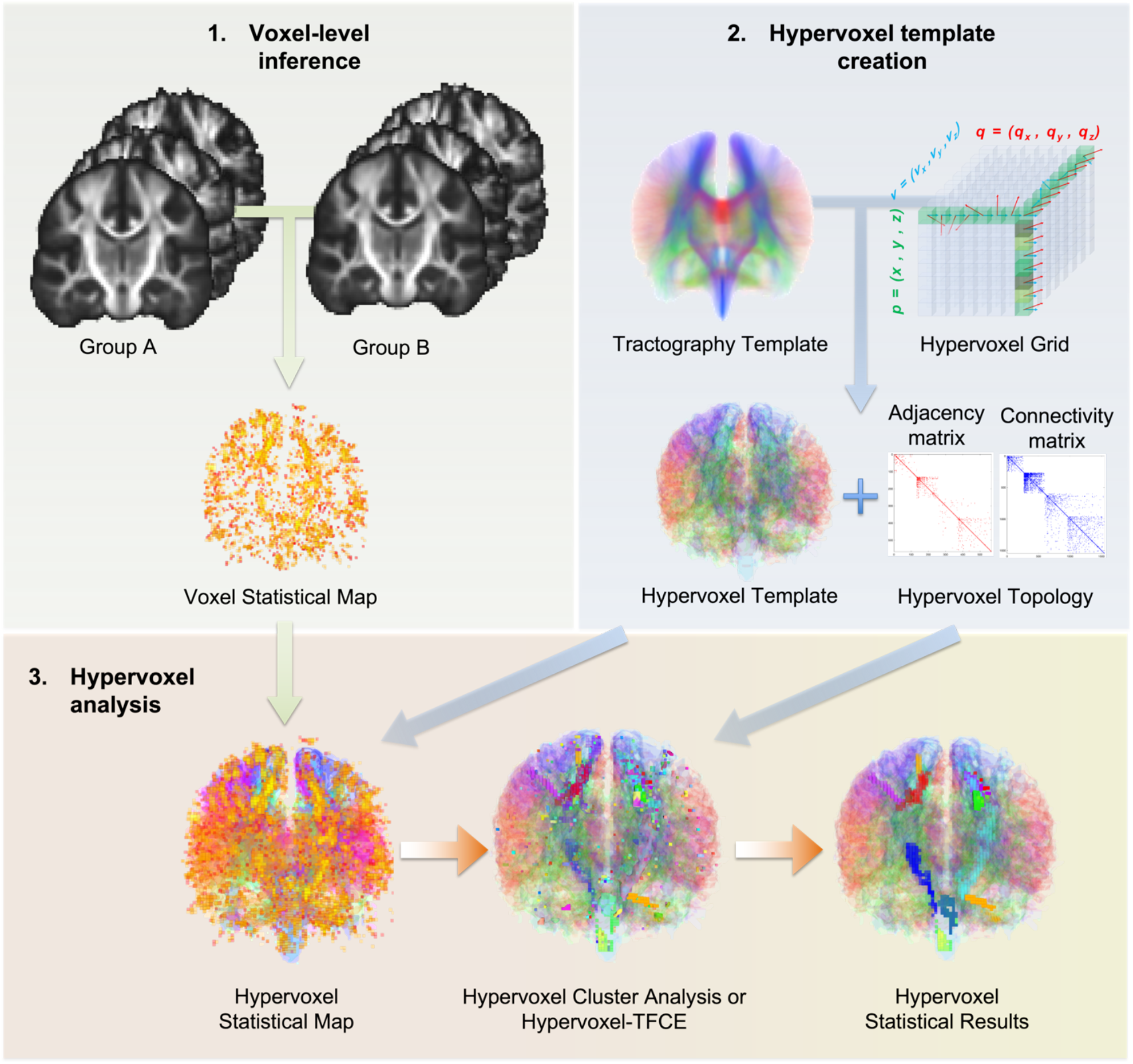
Hypervoxel-based analysis workflow. 1) Statistical maps representing the results of whole-brain voxel-wise statistical inference analysis, for example to detect group level differences between two groups. 2) Hypervoxel template and corresponding connectivity and adjacency matrices obtained from the hypervoxel encoding of tractography data registered to the same of the statistical maps. 3) The statistical maps are projected to the hypervoxel template to be analysed with techniques adapted to the hypervoxel framework such as hypervoxel cluster analysis or hypervoxel TFCE.

**[Fig.7].**
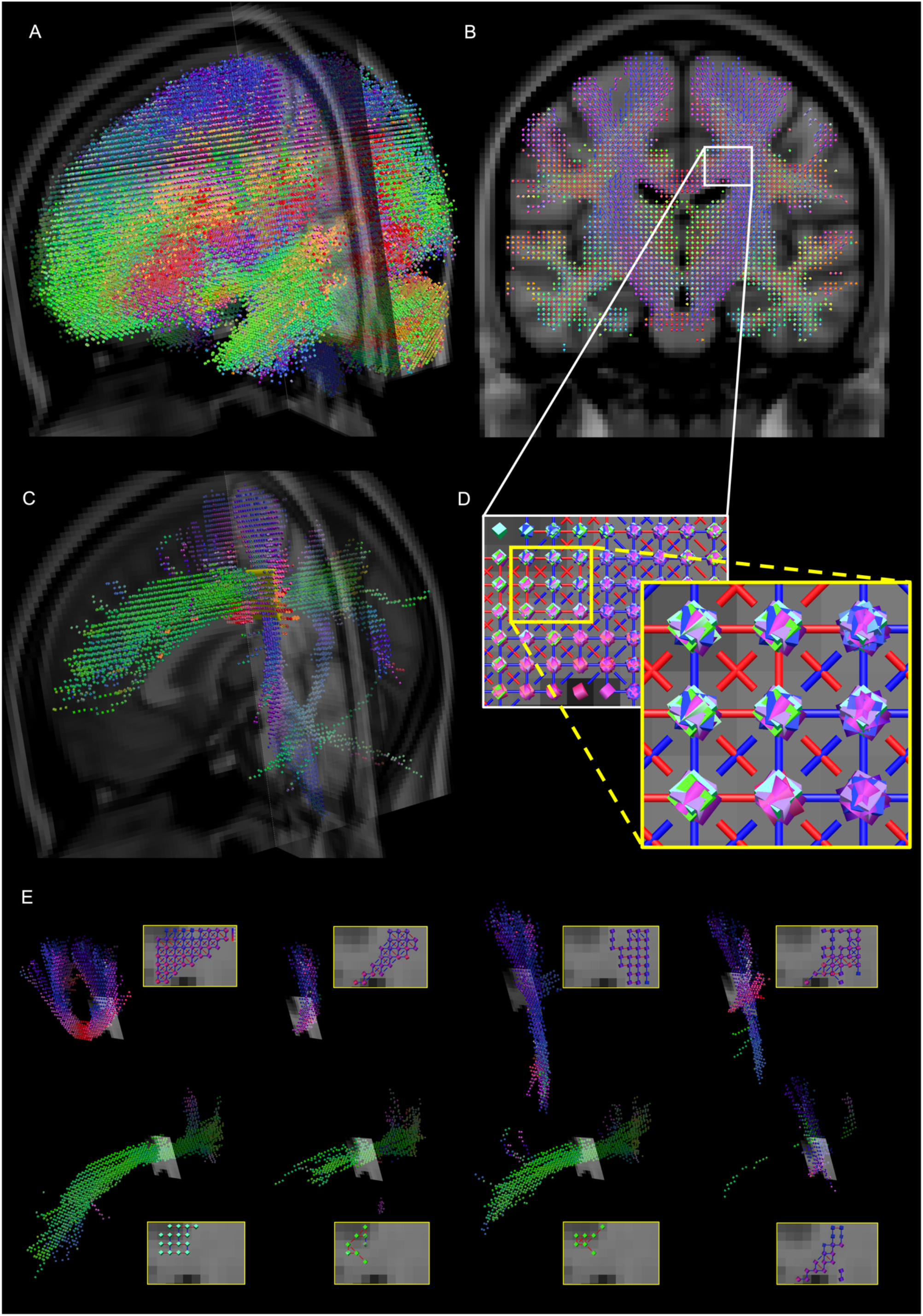
Hypervoxel template. A) 3D visualisation of a whole-brain hypervoxel template where each hypervoxel is represented by an oriented cube with matching RGB colour indicating the local orientation of the hypervoxel. B) Coronal slice of the hypervoxels template at coordinate Y=54 and a delineated 7 × 9 rectangular region within the slide. C) Hypervoxels representing all the streamlines that intersect the 7 × 9 region showed in B. D) Augmented zoom views of the 7 × 9 rectangular region from B showing coloured links indicating the connectivity (blue) and the adjacency (red) between hypervoxels. E) Eight separated connected components obtained from the clustering of the hypervoxels represented in C and D.

### 2.6. Hypervoxel clustering

Based on the hypervoxel topology, we can define a hypervoxel clustering algorithm capable of disentangling overlapping patterns of streamline configurations in regions of “crossing” and “kissing” fibres (Basser et al. 2000). Given a set of voxels intersected by an arbitrary number of tractography streamlines (Fig.5), the objective is to obtain separate clusters of hypervoxels where each cluster represents a differentiated bundle of fibres. This problem cannot be solved in general by conventional voxel-clustering algorithms because each voxel can only be assigned to one cluster, even if the voxel is intersected by multiple fibre populations. Instead, this problem can be overcome by projecting first the voxels into the hypervoxel encoding that represents the tractography data, and then by clustering the resulting subset of hypervoxels using the topological information provided by the connectivity and adjacency matrices. The cluster-growing algorithm iterates through all the hypervoxels in the subset and parsimoniously adds each hypervoxel to an existing cluster if it is either connected or radially adjacent to a hypervoxel already in the cluster.

### 2.7. Hypervoxel template

The hypervoxel encoding of tractography data can be applied separately to individual tractography reconstructions or done simultaneously to multiple tractography datasets registered within the same space. We use the term hypervoxel template to refer to the hypervoxel encoding of tractography data obtained from multiple subjects and normalised to the same 3D image template such as the MNI152 T1 1mm template (Fonov et al. 2009). A hypervoxel template also has associated hypervoxel connectivity and radial adjacency matrices previously defined for its hypervoxel encoding. These matrices provide the information required to perform different topological operations on the hypervoxels, such as the segmentation of a given set of hypervoxels into separated regions of interconnected hypervoxels. To illustrate the topological properties of a hypervoxel template and to demonstrate its applications in neuroimaging, we constructed a hypervoxel template based on whole-brain spherical deconvolution tractography from single-shell diffusion MRI data acquired on 20 healthy adult participants from the BRC-Atlas neuroimaging study on healthy volunteers. These data were collected at KCL Centre for Neuroimaging Science under ethical approval number KCL REC PNM/10/11-163. Each individual tractography dataset was normalised to standard MNI space by applying the same combination of affine and non-linear transformations used to register each of the corresponding FA brain maps to the FMRIB58_FA template available through the FSL software (www.fmrib.ox.ac.uk/fsl) using FSL FLIRT and FNIRT commands (Jenkinson and Smith 2001; Jenkinson et al. 2002, 2012). A series of filters was subsequently applied to the combined tractography data to remove anatomically implausible streamlines including loops, streamlines terminating in the middle of the white matter and streamlines longer than 200mm or shorter than 15mm. In addition, streamlines with trajectories underrepresented across the 20 subjects were filtered down to 100000 streamlines. To reduce the computational demand required to construct the hypervoxel template, we randomly selected a final number of 30000 streamlines using the skip function implemented in Trackvis (Wang and Wedeen 2017).

### 2.8. Hypervoxel cluster analysis and hypervoxel TFCE

Once a hypervoxel template is available, any voxel-based neuroimage data defined in the underlying 3D anatomical space such as the MNI T1 2mm template can be projected to the hypervoxel template to be analysed in hypervoxel space. The projection of data onto the hypervoxel template would generally produce a one-to-many mapping from voxels to hypervoxels covering the entire hypervoxel encoding. In addition, we can apply the previously defined hypervoxel clustering algorithm to the projected data. The strategy of projecting voxel-based data into hypervoxel space before performing a hypervoxel clustering allows us to elegantly extend already existing methods of statistical inference based on voxel clustering also to the hypervoxel framework.

Two examples of cluster based statistical methods that have become standard in the analysis of neuroimaging data are *cluster-wise* inference analysis (Poline and Mazoyer 1993) and threshold-free cluster enhancement (TFCE) (S. M. Smith and Nichols 2009). These two related methodologies work on the assumption that after a whole-brain voxel-level statistical inference analysis, large clusters of statistically significant voxels extending beyond the spatial autocorrelation of the noise are likely to reflect the presence of true effects of interest in the images (Poline and Mazoyer 1993). This same assumption can be applied equally to the analysis of voxel data projected onto a different spatial domain where the topology of the alternative space is used to determine the formation of clusters and their statistical properties (Fischl 2012; Lett et al. 2017; D. A. Raffelt et al. 2015; Zhang, Wu, Ning, et al. 2018). In all these approaches, the topology of the new space is used to determine the formation of clusters and their statistical properties. In a similar manner, we use the hypervoxel topology to implement the hypervoxel versions of cluster-wise inference and TFCE for the analysis of voxel data projected onto a hypervoxel template.

The general workflow of a hypervoxel cluster analysis or a hypervoxel TFCE can be subdivided into four stages that resemble a traditional cluster-wise inference or TFCE in voxel space. Firstly, a typical voxel-level analysis is carried out on the original imaging data using permutation tests (Winkler et al. 2014). This generates for each permutation whole-brain statistical maps that represent the results of evaluating a test hypothesis at the level of individual voxels. Secondly, the statistical maps are projected onto a hypervoxel template created following the steps described in the previous sections. Then, the hypervoxel topology is used to produce for each map either clusters of suprathreshold hypervoxels (in the case of hypervoxel cluster analysis) or the cluster support of each hypervoxel (in the case of a hypervoxel TFCE analysis). The clusters of suprathreshold voxels are obtained by applying the hypervoxel clustering algorithm to those hypervoxels that have a value above a selected arbitrary threshold of the statistic. For the TFCE, the cluster support of each hypervoxel is formed by all other hypervoxels with less or equal value of the statistic connected to a neighbourhood of adjacent hypervoxels. Finally, the cluster-level statistic for each cluster or the TFCE value for each hypervoxel are computed for each permutation test, and their statistical significance under the null hypothesis calculated, using for the control of false positives a family-wise error (FWE) correction based on the distribution of maximal values of the statistic or the TFCE across all permutations.

### 2.9. Application to neuroimaging data

To evaluate the hypervoxel framework, we used hypervoxel cluster analysis and hypervoxel TFCE analysis to detect group-level differences in the brain images of patients diagnosed with amyotrophic lateral sclerosis (ALS), a disease characterised by the selective degeneration of motor neurons along the corticospinal tract (CST) although the pathology also extends to other brain areas (Saberi et al. 2015; Al-Chalabi et al. 2012). In addition to the hypervoxel analyses, we carried out traditional voxel cluster and voxel TFCE analyses on the same data to make direct comparisons between the performance of the two versions of each method (i.e. voxels vs hypervoxels). The analysed data consisted on fractional anisotropy (FA) brain maps derived from diffusion MRI data acquired on 24 (5 female) healthy controls (48.0 years ±8.8 years) and 25 (3 female) limb-onset ALS patients (53.1 years ±12.5 years) using a 3T MRI system (GE Medical Systems HDx) at KCL Centre for Neuroimaging Sciences. Ethics approval for the collection and analysis of the data was obtained from The Joint South London and Maudsley and The Institute of Psychiatry NHS Research Ethics Committee (07/H0807/85) and written informed consent was obtained from each participant following the Declaration of Helsinki.

Each diffusion MRI dataset was acquired using a spin-echo echo-planar imaging twice refocused sequence along 60 contiguous axial slices with the following parameters: TE 104.5 ms, (voxel size 2.4 · 2.4 · 2.4 mm, matrix 128 · 128), 32 diffusion-weighted directions (b-value 1350 s/mm2) and 4 non-diffusion-weighted volumes. The acquisition was gated to the cardiac cycle using a digital pulse oximeter placed on participants’ forefinger. Datasets were pre-processed for motion and eddy current distortion correction using ExploreDTI (Leemans et al. 2009) with the corresponding reorientation of the b-matrix (Leemans and Jones 2009). The diffusion tensor at each voxel was estimated using a non-linear least square approach (Jones and Basser 2004) to produce a whole-brain FA map for each subject that was subsequently normalised to a common anatomical space applying non-linear registration as previously described in (S. M. Smith et al. 2006) using the *tbss_2_reg* FSL script and the FSL FMRIB58_FA template.

We carried out each of the four separate analyses with the aim to detect lower FA values in ALS patients with respect to healthy controls, the most common findings reported by the literature (Gabel et al. 2020; Li et al. 2012). All four analyses investigated the effect of clinical status in voxel FA values using a general linear model (GLM) that included age and sex as covariates and employed permutation tests (5000) for the statistical inference. The first and second analyses consisted of the same voxel-level analysis carried out using the SnPM MATLAB toolbox and followed by separate cluster-level inferences in voxel or hypervoxel space. For each case, clusters of voxels or hypervoxels were computed using the same cluster-forming suprathreshold (equivalent to a p-value = 0.01) followed by the calculation of their cluster-mass (Bullmore et al. 1999). In the case of hypervoxels, we used a modified version of the SnPM toolbox to run the *hypervoxel* cluster analysis based on the provided hypervoxel template. The third and fourth analyses consisted respectively on a voxel and a hypervoxel TFCE analysis of the same FA voxel data. In both cases, the “E”, “H” and neighbourhood-connectivity parameters used for the TFCE calculations were set to the recommended values (*E=0*.*5* and *H=2*) (S. M. Smith and Nichols 2009) using FSL randomise and our modified SnPM MATLAB toolbox for the voxel-TFCE and the hypervoxel TFCE analyses respectively. The results for all four analyses were finally corrected for multiple statistical comparisons using a family-wise error (FWE) p-value = 0.05 based on the distribution of maximal values of the statistic (cluster-level or TFCE) across permutations.

### 2.10. Control for false positives

A requirement for the statistical validity of any inference method is good control for false positives. In a hypervoxel clustering analysis, false positives are expected to occur at a rate no greater than the nominal FWE rate used for the correction of multiple statistical comparisons that controls the probability of reporting one or more false positive results in the analysis. To assess that nominal FWE rates correspond accurately with observed false-positive rates, we empirically measured FWE rates in a hypervoxel cluster analysis by applying the same strategy used by (Eklund, Nichols, and Knutsson 2016) for the detection of group level differences across groups of subjects drawn from a population of healthy controls. In this scenario all positive results are, by definition, false positives. Therefore, the FWE is given by the proportion of analyses that give rise to any significant results. To estimate the FWE rate, we used Fractional Anisotropy (FA) maps based on diffusion MRI data acquired from 200 subjects (50% male) randomly selected from the Human Connectome Project (HCP) (Essen et al. 2012). We retrieved diffusion and T1w images in fully pre-processed form (Glasser et al. 2013), including co-registration and normalisation to the MNI152 T1 1mm template, from the HCP database. We used StarTrack (www.mr-startrack.com) to compute the diffusion tensor and produce FA maps based on the b = 2000 s/mm2 diffusion images. We then applied the affine and warp transformations provided by the HCP to these FA maps, bringing them into the space of the MNI template. We split the 200 FA maps into two random groups of equal size (50% male each) and ran hypervoxel clustering analysis to detect group-level differences in FA. Clusters of supra-threshold hypervoxels (*FA*_*Group*-1_ < *FA*_*Group*-2_, p-value ≤ 0.01 uncorrected) were considered statistically significant if their cluster mass demonstrated an associated FWE p-value < 0.05. We repeated this process 1000 times using different random permutations of the group labels for each run. We compared the empirical FWE rate with its predicted 5% value by plotting the cumulative curve of runs with statistically significant results against the total number of runs.

## 3. Results

### 3.1. Hypervoxel Template

The encoding of the hypervoxel template resulted in approximately 1.5 million encoded hypervoxels representing over 30000 streamlines intersecting around 135000 voxels (Fig.7-A). The distribution of the hypervoxel density in terms of number of hypervoxels per voxel follows a power law distribution where 21% of voxels have a hypervoxel density equal or less than 2, 58% of voxels have a density between 3 and 16, and 21% of voxels have more than 16 hypervoxels per voxel. Voxels with higher hypervoxel density could reflect regions with high number of crossing fibres such as those shown in NuFO maps depicting the number of distinct fibre orientations in each voxel (Descoteaux et al. 2009; F. Dell’Acqua et al. 2013), but they could also indicate the presence of fanning fibres or local parallel fibres that diverge outside the voxel. The overlap of multiple hypervoxels at the same voxel location can be appreciated in a sectional grid of the hypervoxel template depicted as a 3D slice (Fig.7-B) and in the magnified view displaying a selected rectangular region of the same 3D slide (Fig.7-D). All the streamlines that intersect this rectangular region are represented by the hypervoxels depicted in (Fig.7-C). In all these figures, each hypervoxel is depicted by an oriented cube RGB coloured according to its local orientation. However, in the hypervoxel template each hypervoxel is usually disconnected from other hypervoxels occupying the same voxel position. Instead, each hypervoxel can be either connected or adjacent to neighbouring hypervoxels as determined by the hypervoxel connectivity and adjacency matrices. These two relationships allow us to calculate for each hypervoxel an associated region of inter-connected and adjacent hypervoxels, which allow us to “disentangle” the overlapping hypervoxels from (Fig.7-C) and (Fig.7-D) into separate regions illustrated by (Fig.7-E). The topological relationship between hypervoxels within the same “connected” region is now better appreciated by the colour coding of the edges in the template: blue edges indicate the connectivity between neighbouring hypervoxels while red edges indicate their radial adjacency.

### 3.2. Control of false positives

From a total 1000 random hypervoxel cluster analysis tests aimed at detecting group-level differences in FA, only 44 tests reported false-positives in the form of one or more clusters with a statistically significant cluster mass. The empirical false-positive rate associated with the battery of tests follows the predicted 5% FWE rate and remains mostly below it (Fig.8).

**[Fig.8].**
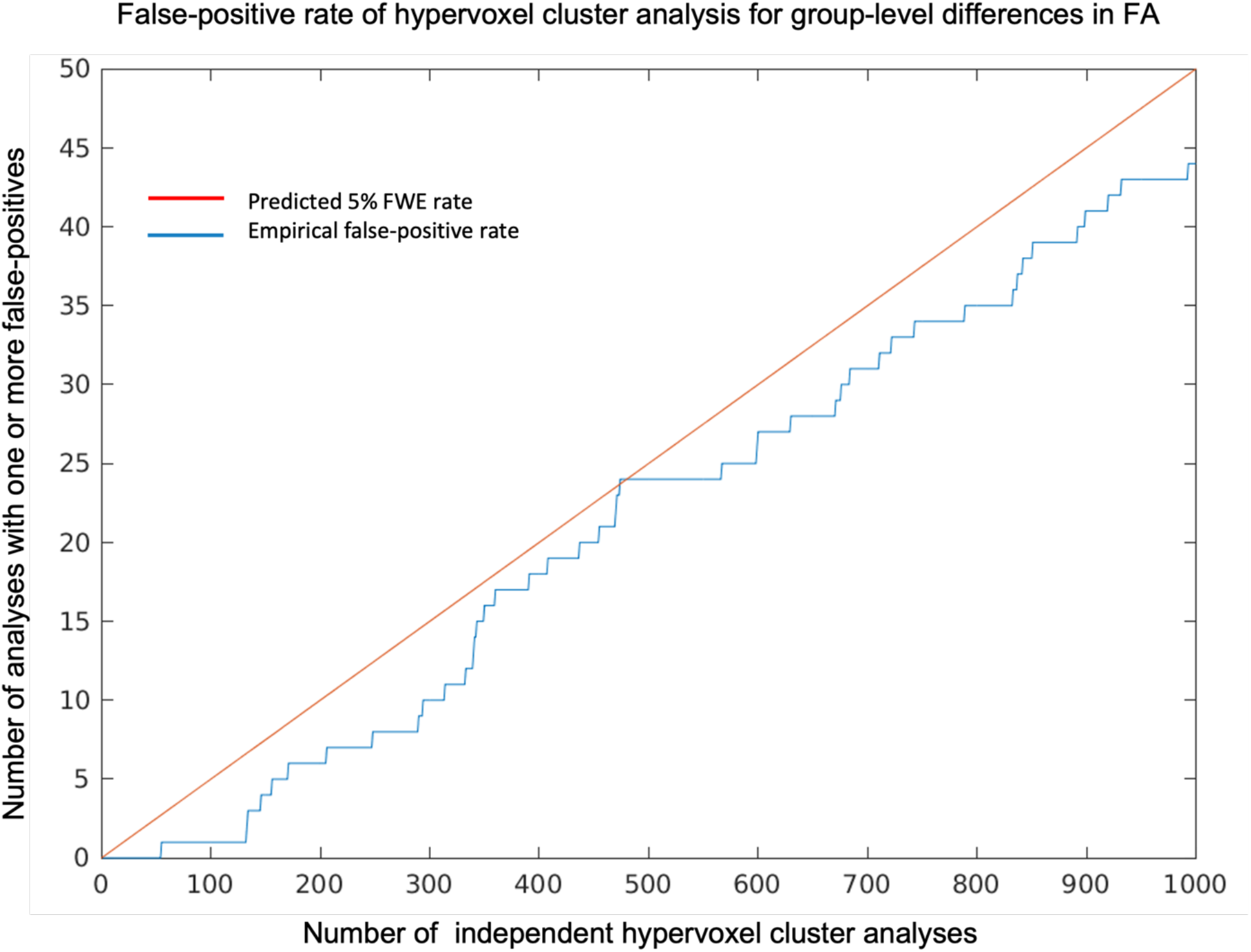
Empirical false-positive rate of a hypervoxel cluster analysis applied to the detection of group level differences in Fractional Anisotropy (FA) between random groups of subjects from the same normal population. Results based on 1000 independent random analysis tests, each one based on a different random split of 200 HCP subjects into two different groups of equal sizes. Each false positive represents one analysis reporting one or more statistically significant clusters of suprathreshold hypervoxels (suprathreshold uncorrected p-value < 0.01, cluster-mass FWE-corrected p-value = 0.05). The empirical false-positive rate remains robustly around or under the predicted 5% FWE rate.

### 3.3. Voxel cluster analysis vs hypervoxel cluster analysis

The results from the *voxel* cluster analysis show two large separate clusters of adjacent suprathreshold voxels where FA is lower in ALS than in healthy controls (Fig.9-Left). One cluster is aligned with part of the inferior branch of the right CST, while the other cluster partially overlaps the superior branch of the left CST and other nearby white matter regions such as the Corpus Callosum (CC) or the left Superior Longitudinal Fascicle (SLF-I). In comparison, the results from the hypervoxel analysis identify a higher number of white matter regions where the FA is significantly lower in ALS (Fig.9-Centre and Right). In total, the cluster-level analysis found 19 clusters of interconnected suprathreshold hypervoxels (*FA*_*ALS*_ < *FA*_*HC*_, p-value ≤ 0.01 uncorrected) with statistically significant cluster mass (p-value ≤ 0.05 *FWE*-corrected). Each cluster lay along one of five major white matter pathways, based on the trajectory of the underlying streamlines (Fig.9-Center): the right-anterior corticospinal tract (CST) (9 clusters, 0.006 ≤ p-value ≤ 0.025), right-posterior CST (1 cluster, p-value = 0.029), left-anterior CST (6 clusters, 0.003 ≤ p-value ≤ 0.042), left-posterior CST (2 clusters, p-values = (0.006,0.048)) and the CC (1 cluster, p-value = 0.003).

**[Fig.9].**
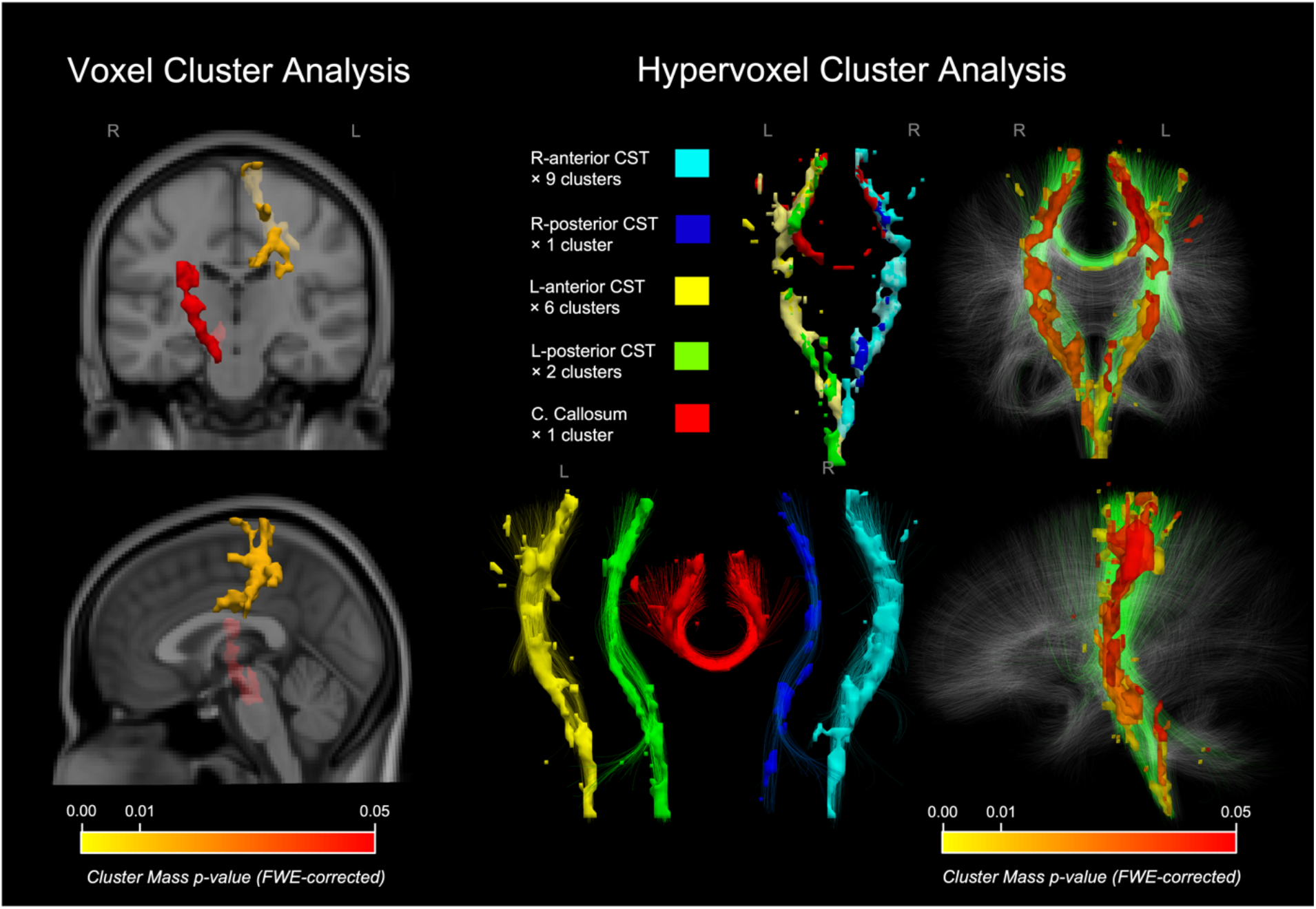
Voxel vs Hypervoxel Cluster Analysis of reduction in fractional anisotropy (FA) in ALS. Left) Statistically significant clusters of voxels colour-coded by p-value (cluster mass). Centre) Statistically significant clusters of hypervoxels categorised into five white-matter tracts based on the underlying streamlines: right-anterior corticospinal tract (CST) (9 clusters), right-posterior CST (1 clusters), left-anterior CST (6 clusters), left-posterior CST (2 clusters) and corpus callosum (CC) (1 cluster). Right) Same statistically significant clusters of hypervoxels colour-coded by p-value (cluster mass) and displayed along with underlying streamlines.

### 3.4. Voxel TFCE analysis vs hypervoxel TFCE analysis

The TFCE analyses approximately replicated the results from the cluster analyses but at the spatial resolution of individual voxels and hypervoxels. There were around 500 statistically significant voxels distributed within the same space occupied by the two clusters from the voxel cluster analyses (Fig.10-Left). In comparison, the threshold-free cluster enhancement analysis yielded more than 6,000 significant hypervoxels with p-values between 0.001 and 0.05 (Fig.10-Right). The significant hypervoxels were extensively distributed along the CST and many of them along the part of the CC that interconnects the left and right motor cortices. A smaller number of significant hypervoxels were also found on the part of the SLF-I that overlaps with the left CST and also along the superior cerebellar peduncles.

**[Fig.10].**
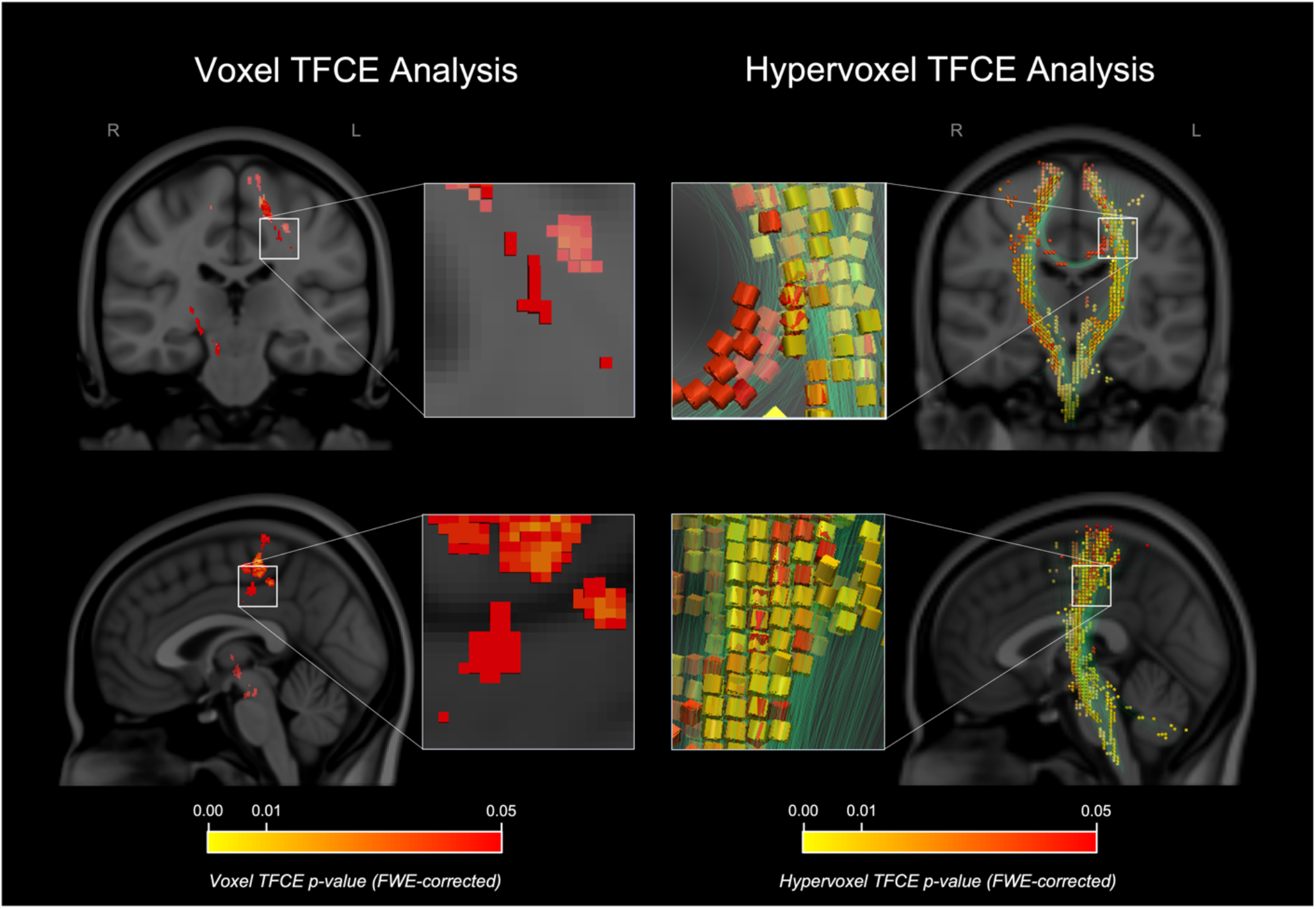
Voxel vs Hypervoxel TFCE Analysis of fractional anisotropy (FA) reduction in ALS. Left) Statistically significant voxels colour-coded by p-value (TFCE). Right) Statistically significant hypervoxels colour-coded by p-value (hypervoxel TFCE) and displayed along the underlying streamlines. Compared to the voxel analysis, the hypervoxels with significant low FA extend also dorsally to the hand region of the precentral gyrus and ventrally along the entire course of the CST, including the fibres of the pyramidal eminences.

## 4. Discussion

The differences observed between the results from the voxel and the hypervoxel analyses reveal the effect that the topological information provided by the hypervoxel template has in the ability of the methods to detect significant effects of interest across the brain. Qualitatively, the results from the hypervoxel analyses can be interpreted with a greater degree of anatomical specificity, both at the scale represented by clusters and the level of individual hypervoxels. Clusters from the hypervoxel cluster analysis appear coherently aligned with the anatomy of specific white-matter tracts, even if the clusters happen to overlap with each other (partially or entirely) in 3D space. This reflects the ability of the method to independently detect separate effects of interest specific to different but overlapping anatomical regions (i.e. white-mater tracts). This is clearly illustrated by the overlapping clusters that correspond separately to callosal and corticospinal pathways, and by the different clusters that partially overlap over sections of the CST. The white-matter pathways associated with each cluster of hypervoxels are determined by the different group of streamlines that the corresponding hypervoxels represent. This anatomical specificity also applies, at the level of individual hypervoxels, to the results from the hypervoxel TFCE analysis. Each statistically significant hypervoxel denotes the presence of an effect of interest specific to the white-matter pathways (i.e. tractography streamlines) locally represented by the hypervoxel. This again represents an increase in “anatomical” resolution with respect to the corresponding voxel analysis that can also be observed by comparing the results from both TFCE analyses. At each voxel location, the hypervoxel TFCE analysis can differentiate between multiple hypervoxels to determine which of them are statistically significant. Such degree of anatomical specificity cannot be achieved, in general, by any voxel-based analysis because the anatomical information provided by voxels is limited to 3D spatial positions and because the topology of the voxel grid is devoid of any anatomical content.

The other noticeable effect is the overall increase in the quantity, statistical significance, and anatomical distribution of the results in the case of the hypervoxel analyses. There were 19 statistically significant hypervoxel clusters (p-value < 0.05,mean p-value = 0.015) compared to only 2 statistically significant voxel clusters (p-value < 0.05,mean p-value = 0.026). Also, there were 6277 significant hypervoxels (p-value < 0.05,mean p-value = 0.0383) compared to only 514 significant voxels (p-value < 0.05,mean p-value = 0.0383) in the TFCE analyses. The fact that the hypervoxel results were more widely spread across the entire CST, the white matter pathway most severely affected in ALS (Foerster et al. 2012; Foerster, Welsh, and Feldman 2013; R. A. L. Menke et al. 2017; Müller et al. 2016), suggests an increased capacity to detect the location of neurodegenerative processes in this disorder, a thesis also supported by the higher statistical significance in the results from the hypervoxel analyses. The robust control for false positives, demonstrated by the empirical estimation of its value at 5% FWE, adds further credibility to the validity of the results including those detected outside the CST. In this respect, the significant results specific to the CC and the SLF agree with previous reports of differences in diffusion MRI metrics in that area in ALS (Filippini et al. 2010; Gabel et al. 2020; D. A. Raffelt et al. 2015; Ricarda A. L. Menke et al. 2014). The results located towards the cerebellum also concur with reported FA reductions in cerebro-cerebellar tracts and the cerebellar peduncles based on a much larger sample of ALS patients (Bede et al. 2021). This apparent increase in “sensitivity” should be mostly attributed to the effect that the topological and anatomical information provided by the hypervoxel encoding has on the statistical inference, as everything else was kept fundamentally the same across the different analyses. The statistical significance of each result (either cluster or TFCE) depends on the magnitude of its cluster-level statistic (under the null hypothesis) with respect to the distribution (across permutation tests) of maximal values of the statistic over all regions (either clusters or TFCE supports). Therefore, the effectiveness of the analysis depends on its capacity to automatically select regions where the values of the final statistic are consistently higher under the null hypothesis than across random permutations of the data. In a hypervoxel analysis this is precisely the case when the effects of interest in the neuroimaging data are topologically aligned with the anatomy represented by the tractography streamlines. In the analysis of the FA images in ALS data this hypothesis holds true to a high degree, hence the observed increase in the statistical significance of the results with respect to the voxel-based analyses.

The tractography data used to create the hypervoxel template should be regarded as anatomical hypotheses introduced a priori by the hypervoxel framework for the analysis of the neuroimaging data. In this respect, there are no general restrictions for the tractography data other than the final hypervoxel template should be relevant to the research hypothesis. For example, a study focused only on short-range connections could use tractography data containing only u-shape fibres. In this study we used whole-brain tractography data to demonstrate the ability of the method to detect effects of interest across the entire white matter. Had we been interested in detecting differences along the CST and the CC only, we could have used tractography data containing only those tracts to boost the statistical power in the same fashion of a region of interest (ROI) analysis. Another factor to consider is the population from which the tractography data are obtained. We used high-quality tractography data acquired on 20 healthy individuals from another study (Howells et al. 2018) to show that a hypervoxel analysis does not require tractography data from the same subjects whose images are analysed. This allows the framework to be applied in studies where no tractography or even no diffusion data are available and facilitates the comparison of results across studies where the same hypervoxel template is used for the analysis. This should not prevent the use of study-specific hypervoxel templates when deemed necessary, for example when the anatomical variability of the population of interest deviates significantly from that of the normal population. A last factor to consider is that the presence of tractography artefacts could increase the likelihood of finding anatomically implausible results (Maier-Hein et al. 2017; C.-H. Yeh et al. 2021), therefore the use of quality tractography data is key. We used comprehensive anatomical filters to remove artefacts from the tractography data at the risk of also removing streamlines representing genuine anatomical connections. This trade-off will remain until specific criteria that confirm the validity of the anatomical connections represented by the streamlines become available. In this respect, the hypervoxel framework will benefit from any future improvements in the field of tractography and at the very least, the number of false-positive results in a hypervoxel analysis (anatomically plausible *and* implausible) will always remain below the chosen statistical significance level for the analysis (i.e. the *p-value*). Above all, the framework provides users the flexibility to create a hypervoxel template based on their own choice of tractography data such as third-party tractography atlases (Catani and Schotten 2008; Hansen et al. 2021; Román et al. 2017; F.-C. Yeh et al. 2018; Zhang, Wu, Norton, et al. 2018) or by using the latest tractography methods available in the field.

In this study, we use the hypervoxel framework only for the final analysis of the statistical maps that resulted from the voxel-level analyses. This approach does not utilise all the potential of the hypervoxels for dealing with the limitations that arise from the interindividual variability found in the white-matter anatomy (Flavio Dell’Acqua and Catani 2012). In a voxel-level analysis, the same voxel in common space can represent very different tracts across subjects. This problem cannot be solved by spatial normalisation of the 3D images alone because the topological embedding of the neuronal connections can be very different from subject to subject and even entire tracts can be missed in some subjects. However, this situation could be greatly ameliorated by starting the analysis of the image data directly in the hypervoxel space defined for each individual, which would be the scope of future works. This approach will be particularly pertinent for the analysis of the apparent fibre density (AFD) (D. Raffelt et al. 2012) and the Hindrance Modulated Orientational Anisotropy (HMOA) (F. Dell’Acqua et al. 2013) given by the amplitude of the lobes of the fibre ODF obtained through the spherical deconvolution of the diffusion MRI signal (F. Dell’Acqua et al. 2007, 2010; Tournier et al. 2004; Tournier, Calamante, and Connelly 2007). Also promising is the prospective analysis of tract-specific measurements provided by methods that use tractography data to predict at the voxel level the signal from diffusion MRI (Daducci et al. 2015; R. E. Smith et al. 2015; S. Schiavi et al. 2020) and other image modalities (S. Schiavi et al. 2022). The high specificity of the hypervoxels with respect to the streamlines from which these measurements are derived should made of them an ideal framework for the analysis of this kind of data.

## 5. Conclusions

In this work, we introduce hypervoxels as a new methodological framework that combines the encoding capabilities of multidimensional voxels with anatomical and topological information provided by tractography data. By extending the 3D voxel grid with new dimensions to encode information about the streamlines, the new framework reconciles the “voxel space” of the 3D images with the “streamline space” defined by diffusion MRI tractography reconstructions. The added dimensions increase the anatomical specificity of the hypervoxel grid and allow the definition of a new hypervoxel topology that better reflects the structural connectivity of the brain as represented by the tractography streamlines. The use of hypervoxels in the analysis of diffusion MRI data from a study on ALS was associated with improved ability to detect significant effects of interest in the images and greater anatomical accuracy in the results. The apparent increase in sensitivity and specificity can be explained by the greater ability of the hypervoxels to represent the anatomical and topological complexity of the brain’s white matter. We expect that the adoption of a hypervoxel methodology would improve the performance of neuroimaging analyses, especially when investigating phenomena that manifest in the images with a high degree of spatial correlation with respect to the white matter anatomy.

## 7. Acknowledgements

Data were provided [in part] by the Human Connectome Project, WU-Minn Consortium (Principal Investigators: David Van Essen and Kamil Ugurbil; 1U54MH091657) funded by the 16 NIH Institutes and Centers that support the NIH Blueprint for Neuroscience Research; and by the McDonnell Center for Systems Neuroscience at Washington University. The authors acknowledge use of the research computing facility at King’s College London, *Rosalind* (https://rosalind.kcl.ac.uk), which is delivered in partnership with the National Institute for Health Research (NIHR) Biomedical Research Centres at South London & Maudsley and Guy’s & St. Thomas’ NHS Foundation Trusts, and part-funded by capital equipment grants from the Maudsley Charity (award 980) and Guy’s & St. Thomas’ Charity (TR130505). The views expressed are those of the author(s) and not necessarily those of the NHS, the NIHR, King’s College London, or the Department of Health and Social Care. Thank you to Jose Luis Jaramillo for his comments on differential geometry, to Cedric Ginestet for early discussions about the idea of hypervoxels, and to Luis Lacerda for proof reading the final manuscript.

## 8. Author Contributions

Hypervoxel design and conception: PLL and FDA. Implementation: PLL. Neuroimaging data: LHG, MC and SCRW. Data pre-processing and analysis: PLL, RS, AB, FDSR and FDA. Evaluation of results: PLL, DKJ, LHG, MC, SCRW and FDA. Figures: PLL. Manuscript: PLL and FDA.

## REFERENCES

Al-Chalabi, Ammar, Ashley Jones, Claire Troakes, Andrew King, Safa Al-Sarraj, and Leonard H Van Den Berg. 2012. “The Genetics and Neuropathology of Amyotrophic Lateral Sclerosis.” Acta Neuropathologica. https://doi.org/10.1007/s00401-012-1022-4.

Alpert, N. M., J. F. Bradshaw, D. Kennedy, and J. A. Correia. 1990. “The Principal Axes Transformation-a Method for Image Registration.” Journal of Nuclear Medicine 31: 1717–22.

Ashburner, John. 2012. “SPM: A History.” NeuroImage 62 (August): 791–800. https://doi.org/10.1016/j.neuroimage.2011.10.025.

Bach, M, F B Laun, A Leemans, C M W Tax, G J Biessels, B Stieltjes, and K H Maier-Hein. 2014. “Methodological Considerations on Tract-Based Spatial Statistics (TBSS).” NeuroImage 100: 358–69. https://doi.org/10.1016/j.neuroimage.2014.06.021.

Basser, Peter J., Sinisa Pajevic, Carlo Pierpaoli, Jeffrey Duda, and Akram Aldroubi. 2000. “In Vivo Fiber Tractography Using DT-MRI Data.” Magnetic Resonance in Medicine 44 (October): 625–32. https://doi.org/10.1002/1522-2594(200010)44:4%3C625::AID-MRM17%3E3.0.CO;2-O.

Bassett, Danielle S, Daniel L Greenfield, Andreas Meyer-Lindenberg, Daniel R Weinberger, Simon W Moore, and Edward T Bullmore. 2010. “Efficient Physical Embedding of Topologically Complex Information Processing Networks in Brains and Computer Circuits.” PLOS Computational Biology 6 (April): e1000748.#x2013;. https://doi.org/10.1371/journal.pcbi.1000748.

Bede, Peter, Rangariroyashe H. Chipika, Foteini Christidi, Jennifer C. Hengeveld, Efstratios Karavasilis, Georgios D. Argyropoulos, Jasmin Lope, et al. 2021. “Genotype-Associated Cerebellar Profiles in ALS: Focal Cerebellar Pathology and Cerebro-Cerebellar Connectivity Alterations.” Journal of Neurology, Neurosurgery & Psychiatry 92 (November): 1197–1205. https://doi.org/10.1136/jnnp-2021-326854.

Bell, S., M. Cercignani, S. Deoni, Y. Assaf, O. Pasternak, J. Evans, A. Leemans, and D K Jones. 2011. “Ismrm2011.” In Tractometry - Comprehensive Multi-Modal Quantitative Assessment of White Matter Along Specific Tracts, Abstract 0678.

Bullmore, E T, J Suckling, S Overmeyer, S Rabe-Hesketh, E Taylor, and M J Brammer. 1999. “Global, Voxel, and Cluster Tests, by Theory and Permutation, for a Difference Between Two Groups of Structural MR Images of the Brain.” IEEE Transactions on Medical Imaging 18: 32–42. https://doi.org/10.1109/42.750253.

Castro, I De, D D H Christoph, D P Dos Santos, and J A Landeiro. 2005. “Internal Structure of the Cerebral Hemispheres: An Introduction of Fiber Dissection Technique.” Arquivos de Neuro-Psiquiatria 63: 252–58. https://doi.org/10.1590/s0004-282x2005000200011.

Catani, M, R J Howard, S Pajevic, and D K Jones. 2002. “Virtual in Vivo Interactive Dissection of White Matter Fasciculi in the Human Brain.” NeuroImage 17: 77–94. https://doi.org/10.1006/nimg.2002.1136.

Catani, M, and M Thiebaut de Schotten. 2008. “A Diffusion Tensor Imaging Tractography Atlas for Virtual in Vivo Dissections.” Cortex 44 (September): 1105–32. https://doi.org/10.1016/j.cortex.2008.05.004.

Cox, Robert W. 2012. “AFNI: What a Long Strange Trip It’s Been.” NeuroImage 62 (August): 743–47. https://doi.org/10.1016/j.neuroimage.2011.08.056.

Daducci, Alessandro, Alessandro Dal Palu, Alia Lemkaddem, and Jean-Philippe Thiran. 2015. “COMMIT: Convex Optimization Modeling for Microstructure Informed Tractography.” IEEE Transactions on Medical Imaging 34 (January): 246–57. https://doi.org/10.1109/TMI.2014.2352414.

Dell’Acqua, Flavio, and Marco Catani. 2012. “Structural Human Brain Networks.” Current Opinion in Neurology, July, 1. https://doi.org/10.1097/WCO.0b013e328355d544.

Dell’Acqua, F., G. Rizzo, P. Scifo, R. A. Clarke, G. Scotti, and F. Fazio. 2007. “A Model-Based Deconvolution Approach to Solve Fiber Crossing in Diffusion-Weighted MR Imaging.” IEEE Transactions on Biomedical Engineering 54: 462–72. https://doi.org/10.1109/TBME.2006.888830.

Dell’Acqua, F., P. Scifo, G. Rizzo, M. Catani, A. Simmons, G. Scotti, and F. Fazio. 2010. “A Modified Damped Richardson-Lucy Algorithm to Reduce Isotropic Background Effects in Spherical Deconvolution.” NeuroImage 49: 1446–58. https://doi.org/10.1016/j.neuroimage.2009.09.033.

Dell’Acqua, F., A. Simmons, S. C. R. Williams, and M. Catani. 2013. “Can Spherical Deconvolution Provide More Information Than Fiber Orientations? Hindrance Modulated Orientational Anisotropy, a True-Tract Specific Index to Characterize White Matter Diffusion.” Human Brain Mapping 34: 2464–83. https://doi.org/10.1002/hbm.22080.

Dell’Acqua, F., and J.-D. Tournier. 2019. “Modelling White Matter with Spherical Deconvolution: How and Why?” NMR in Biomedicine 32. https://doi.org/10.1002/nbm.3945.

Descoteaux, M., R. Deriche, T. R. Knösche, and A. Anwander. 2009. “Deterministic and Probabilistic Tractography Based on Complex Fibre Orientation Distributions.” IEEE Transactions on Medical Imaging 28: 269–86. https://doi.org/10.1109/TMI.2008.2004424.

Ehresmann, Charles. 1950. “Les Connexions Infinitésimales Dans Un Espace Fibré Différentiable.” Séminaire Bourbaki 1: 153–68. http://eudml.org/doc/109403.

Eklund, Anders, Thomas E Nichols, and Hans Knutsson. 2016. “Cluster Failure: Why fMRI Inferences for Spatial Extent Have Inflated False-Positive Rates.” Proceedings of the National Academy of Sciences 113 (July): 7900–7905. http://www.pnas.org/content/113/28/7900.abstract.

Essen, D C Van, K Ugurbil, E Auerbach, D Barch, T E J Behrens, R Bucholz, A Chang, et al. 2012. “The Human Connectome Project: A Data Acquisition Perspective.” NeuroImage 62 (October): 2222–31. https://doi.org/http://dx.doi.org/10.1016/j.neuroimage.2012.02.018.

Farquharson, S., J.-D. Tournier, F. Calamante, G. Fabinyi, M. Schneider-Kolsky, G. D. Jackson, and A. Connelly. 2013. “White Matter Fiber Tractography: Why We Need to Move Beyond DTI.” Journal of Neurosurgery 118: 1367–77. https://doi.org/10.3171/2013.2.JNS121294.

Filippini, N., G. Douaud, C. E. MacKay, S. Knight, K. Talbot, and M. R. Turner. 2010. “Corpus Callosum Involvement Is a Consistent Feature of Amyotrophic Lateral Sclerosis.” Neurology 75: 1645–52. https://doi.org/10.1212/WNL.0b013e3181fb84d1.

Fischl, Bruce. 2012. “FreeSurfer.” NeuroImage 62 (August): 774–81. https://doi.org/10.1016/J.NEUROIMAGE.2012.01.021.

Foerster, B. R., B. A. Dwamena, M. Petrou, R. C. Carlos, B. C. Callaghan, and M. G. Pomper. 2012. “Diagnostic Accuracy Using Diffusion Tensor Imaging in the Diagnosis of ALS. A Meta-Analysis.” Academic Radiology 19: 1075–86. https://doi.org/10.1016/j.acra.2012.04.012.

Foerster, B. R., R. C. Welsh, and E. L. Feldman. 2013. “25 Years of Neuroimaging in Amyotrophic Lateral Sclerosis.” Nature Reviews Neurology 9: 513–24. https://doi.org/10.1038/nrneurol.2013.153.

Fonov, VS, AC Evans, RC McKinstry, CR Almli, and DL Collins. 2009. “Unbiased Nonlinear Average Age-Appropriate Brain Templates from Birth to Adulthood.” NeuroImage 47 (July): S102. https://doi.org/10.1016/S1053-8119(09)70884-5.

Fornito, A., A. Zalesky, and M. Breakspear. 2015. “The Connectomics of Brain Disorders.” Nature Reviews Neuroscience 16: 159–72. https://doi.org/10.1038/nrn3901.

Friston, K J, J Ashburner, C D Frith, J B Poline, J D Heather, and R S J Frackowiak. 1995. “Spatial Registration and Normalization of Images.” Human Brain Mapping 3: 165–89. http://www.scopus.com/inward/record.url?eid=2-s2.0-0029197929&partnerID=40&md5=5a5b553b557d7ef8abac5741af7082bb.

Friston, K. J., P. Jezzard, and R. Turner. 1994. “Analysis of Functional MRI Time-series.” Human Brain Mapping 1: 153–71. https://doi.org/10.1002/hbm.460010207.

Gabel, M. C., R. J. Broad, A. L. Young, S. Abrahams, M. E. Bastin, R. A. L. Menke, A. Al-Chalabi, et al. 2020. “Evolution of White Matter Damage in Amyotrophic Lateral Sclerosis.” Annals of Clinical and Translational Neurology 7: 722–32. https://doi.org/10.1002/acn3.51035.

Glasser, Matthew F, Stamatios N Sotiropoulos, J A Wilson, T S Coalson, Bruce Fischl, J L Andersson, J Xu, et al. 2013. “The Minimal Preprocessing Pipelines for the Human Connectome Project.” NeuroImage 80 (October): 105–24. https://doi.org/10.1016/j.neuroimage.2013.04.127.

Hagmann, Patric. 2005. “From Diffusion MRI to Brain Connectomics.” EPFL.

Hagmann, Patric, Leila Cammoun, Xavier Gigandet, Stephan Gerhard, P. Ellen Grant, Van Wedeen, Reto Meuli, Jean-Philippe Thiran, Christopher J. Honey, and Olaf Sporns. 2010. “MR Connectomics: Principles and Challenges.” Journal of Neuroscience Methods 194 (December): 34–45. https://doi.org/10.1016/j.jneumeth.2010.01.014.

Hansen, Colin B, Qi Yang, Ilwoo Lyu, Francois Rheault, Cailey Kerley, Bramsh Qamar Chandio, Shreyas Fadnavis, et al. 2021. “Pandora: 4-d White Matter Bundle Population-Based Atlases Derived from Diffusion MRI Fiber Tractography.” Neuroinformatics 19 (July): 447–60. https://doi.org/10.1007/s12021-020-09497-1.

Howells, H, M T De Schotten, F Dell’Acqua, A Beyh, G Zappalà, A Leslie, A Simmons, D G Murphy, and M Catani. 2018. “Frontoparietal Tracts Linked to Lateralized Hand Preference and Manual Specialization.” Cerebral Cortex 28. https://doi.org/10.1093/cercor/bhy040.

Ivancevic, V G, and T T Ivancevic. 2007. Applied Differential Geometry: A Modern Introduction. Applied Differential Geometry: A Modern Introduction. https://doi.org/10.1142/6420.

Jenkinson, M, P Bannister, M Brady, and S M Smith. 2002. “Improved Optimization for the Robust and Accurate Linear Registration and Motion Correction of Brain Images.” NeuroImage 17: 825–41. https://doi.org/10.1016/S1053-8119(02)91132-8.

Jenkinson, M, C Beckmann, T E Behrens, M Woolrich, and S M Smith. 2012. “FSL.” NeuroImage 62 (August): 782–90. https://doi.org/10.1016/J.NEUROIMAGE.2011.09.015.

Jenkinson, M, and S M Smith. 2001. “A Global Optimisation Method for Robust Affine Registration of Brain Images.” Medical Image Analysis 5: 143–56. https://doi.org/10.1016/S1361-8415(01)00036-6.

Jeurissen, B., M. Descoteaux, S. Mori, and A. Leemans. 2019. “Diffusion MRI Fiber Tractography of the Brain.” NMR in Biomedicine 32. https://doi.org/10.1002/nbm.3785.

Johansen-Berg, H., and M. F. S. Rushworth. 2009. Using Diffusion Imaging to Study Human Connectional Anatomy. Annual Review of Neuroscience. Vol. 32. https://doi.org/10.1146/annurev.neuro.051508.135735.

Jones, D K, and P J Basser. 2004. “”Squashing Peanuts and Smashing Pumpkins”: How Noise Distorts Diffusion-Weighted MR Data.” Magnetic Resonance in Medicine 52: 979–93. https://doi.org/10.1002/mrm.20283.

Jones, D K, and M Nilsson. 2015. “Microstructures of Learning: Novel Methods and Approaches for Assessing Structural and Functional Changes Underlying Knowledge Acquisition in the Brain.” In Tractometry and the Hunt for the Missing Link: A Physicist Perspective, edited by Merle Horne, 38–48. Frontiers Media SA. https://doi.org/10.3389/978-2-88919-480-3.

Kolář, Ivan, Jan Slovák, and Peter W. Michor. 1993. Natural Operations in Differential Geometry. Springer Berlin Heidelberg. https://doi.org/10.1007/978-3-662-02950-3.

Leemans, A, B Jeurissen, J Sijbers, and D K Jones. 2009. “ExploreDTI: A Graphical Toolbox for Processing, Analyzing, and Visualizing Diffusion MR Data.” 17th Annual Meeting of Intl Soc Mag Reson Med.

Leemans, A, and D K Jones. 2009. “The b-Matrix Must Be Rotated When Correcting for Subject Motion in DTI Data.” Magnetic Resonance in Medicine 61: 1336–49. https://doi.org/10.1002/mrm.21890.

Lett, Tristram A., Lea Waller, Heike Tost, Ilya M. Veer, Arash Nazeri, Susanne Erk, Eva J. Brandl, et al. 2017. “Cortical Surface-Based Threshold-Free Cluster Enhancement and Cortexwise Mediation.” Human Brain Mapping 38 (June): 2795–2807. https://doi.org/10.1002/hbm.23563.

Li, JianPeng, PingLei Pan, Wei Song, Rui Huang, Ke Chen, and HuiFang Shang. 2012. “A Meta-Analysis of Diffusion Tensor Imaging Studies in Amyotrophic Lateral Sclerosis.” Neurobiology of Aging 33 (August): 1833–38. https://doi.org/10.1016/j.neurobiolaging.2011.04.007.

Maier-Hein, K. H., P. F. Neher, J.-C. Houde, M.-A. Côté, E. Garyfallidis, J. Zhong, M. Chamberland, et al. 2017. “The Challenge of Mapping the Human Connectome Based on Diffusion Tractography.” Nature Communications 8. https://doi.org/10.1038/s41467-017-01285-x.

Menke, R. A. L., F. Agosta, J. Grosskreutz, M. Filippi, and M. R. Turner. 2017. “Neuroimaging Endpoints in Amyotrophic Lateral Sclerosis.” Neurotherapeutics 14: 11–23. https://doi.org/10.1007/s13311-016-0484-9.

Menke, Ricarda A. L., Sonja Körner, Nicola Filippini, Gwenaëlle Douaud, Steven Knight, Kevin Talbot, and Martin R. Turner. 2014. “Widespread Grey Matter Pathology Dominates the Longitudinal Cerebral MRI and Clinical Landscape of Amyotrophic Lateral Sclerosis.” Brain 137 (September): 2546–55. https://doi.org/10.1093/brain/awu162.

Müller, H.-P., M. R. Turner, J. Grosskreutz, S. Abrahams, P. Bede, V. Govind, J. Prudlo, A. C. Ludolph, M. Filippi, and J. Kassubek. 2016. “A Large-Scale Multicentre Cerebral Diffusion Tensor Imaging Study in Amyotrophic Lateral Sclerosis.” Journal of Neurology, Neurosurgery and Psychiatry 87: 570–79. https://doi.org/10.1136/jnnp-2015-311952.

Nimsky, C., O. Ganslandt, and R. Fahlbusch. 2006. “Implementation of Fiber Tract Navigation.” Neurosurgery 58. https://doi.org/10.1227/01.NEU.0000204726.00088.6D.

Pineda-Pardo, José Ángel, Kenia Martínez, Ana Beatriz Solana, Juan Antonio Hernández-Tamames, Roberto Colom, and Francisco del Pozo. 2015. “Disparate Connectivity for Structural and Functional Networks Is Revealed When Physical Location of the Connected Nodes Is Considered.” Brain Topography 28 (March): 187–96. https://doi.org/10.1007/s10548-014-0393-3.

Poline, J.-B., and B M Mazoyer. 1993. “Analysis of Individual Positron Emission Tomography Activation Maps by Detection of High Signal-to-Noise-Ratio Pixel Clusters.” Journal of Cerebral Blood Flow and Metabolism 13: 425–37. https://www.scopus.com/inward/record.uri?eid=2-s2.0-0027287403&partnerID=40&md5=a5b4577a9437e9d3726b4214d971a90e.

Raffelt, David A, Robert E Smith, Gerard R Ridgway, J-Donald Tournier, David N Vaughan, Stephen Rose, Robert Henderson, and Alan Connelly. 2015. “Connectivity-Based Fixel Enhancement: Whole-Brain Statistical Analysis of Diffusion MRI Measures in the Presence of Crossing Fibres.” NeuroImage 117 (August): 40–55. https://doi.org/http://dx.doi.org/10.1016/j.neuroimage.2015.05.039.

Raffelt, D., J.-D. Tournier, S. Rose, G. R. Ridgway, R. Henderson, S. Crozier, O. Salvado, and A. Connelly. 2012. “Apparent Fibre Density: A Novel Measure for the Analysis of Diffusion-Weighted Magnetic Resonance Images.” NeuroImage 59: 3976–94. https://doi.org/10.1016/j.neuroimage.2011.10.045.

Román, Claudio, Miguel Guevara, Ronald Valenzuela, Miguel Figueroa, Josselin Houenou, Delphine Duclap, Cyril Poupon, Jean-François Mangin, and Pamela Guevara. 2017. “Clustering of Whole-Brain White Matter Short Association Bundles Using HARDI Data.” Frontiers in Neuroinformatics 11 (December). https://doi.org/10.3389/fninf.2017.00073.

Saberi, S., J. E. Stauffer, D. J. Schulte, and J. Ravits. 2015. “Neuropathology of Amyotrophic Lateral Sclerosis and Its Variants.” Neurologic Clinics 33: 855–76. https://doi.org/10.1016/j.ncl.2015.07.012.

Schiavi, S, Po-Jui Lu, M Weigel, A Lutti, D K Jones, L Kappos, C Granziera, and A Daducci. 2022. “Bundle Myelin Fraction (BMF) Mapping of Different White Matter Connections Using Microstructure Informed Tractography.” NeuroImage 249 (April): 118922. https://doi.org/10.1016/j.neuroimage.2022.118922.

Schiavi, S., M. Ocampo-Pineda, M. Barakovic, L. Petit, M. Descoteaux, J.-P. Thiran, and A. Daducci. 2020. “A New Method for Accurate in Vivo Mapping of Human Brain Connections Using Microstructural and Anatomical Information.” Science Advances 6. https://doi.org/10.1126/sciadv.aba8245.

Schnepel, P, A Kumar, M Zohar, A Aertsen, and C Boucsein. 2015. “Physiology and Impact of Horizontal Connections in Rat Neocortex.” Cerebral Cortex 25: 3818–35. https://doi.org/10.1093/cercor/bhu265.

Siless, Viviana, Juliet Y. Davidow, Jared Nielsen, Qiuyun Fan, Trey Hedden, Marisa Hollinshead, Elizabeth Beam, et al. 2020. “Registration-Free Analysis of Diffusion MRI Tractography Data Across Subjects Through the Human Lifespan.” NeuroImage 214 (July): 116703. https://doi.org/10.1016/j.neuroimage.2020.116703.

Silva, Susana M, and José Paulo Andrade. 2016. “Neuroanatomy: The Added Value of the Klingler Method.” Annals of Anatomy - Anatomischer Anzeiger 208: 187–93. https://doi.org/https://doi.org/10.1016/j.aanat.2016.06.002.

Smith, R. E., J.-D. Tournier, F. Calamante, and A. Connelly. 2015. “Sift2: Enabling Dense Quantitative Assessment of Brain White Matter Connectivity Using Streamlines Tractography.” NeuroImage 119: 338–51. https://doi.org/10.1016/j.neuroimage.2015.06.092.

Smith, S M, M Jenkinson, H Johansen-Berg, D Rueckert, T E Nichols, C E Mackay, K E Watkins, et al. 2006. “Tract-Based Spatial Statistics: Voxelwise Analysis of Multi-Subject Diffusion Data.” NeuroImage 31: 1487–1505. https://doi.org/10.1016/j.neuroimage.2006.02.024.

Smith, S M, and T E Nichols. 2009. “Threshold-Free Cluster Enhancement: Addressing Problems of Smoothing, Threshold Dependence and Localisation in Cluster Inference.” NeuroImage 44 (January): 83–98. https://doi.org/http://dx.doi.org/10.1016/j.neuroimage.2008.03.061.

Sporns, O., G. Tononi, and R. Kötter. 2005. “The Human Connectome: A Structural Description of the Human Brain.” PLoS Computational Biology 1: 0245–51. https://doi.org/10.1371/journal.pcbi.0010042.

Tournier, J.-D., F. Calamante, and A. Connelly. 2007. “Robust Determination of the Fibre Orientation Distribution in Diffusion MRI: Non-Negativity Constrained Super-Resolved Spherical Deconvolution.” NeuroImage 35: 1459–72. https://doi.org/10.1016/j.neuroimage.2007.02.016.

Tournier, J.-D., F. Calamante, D. G. Gadian, and A. Connelly. 2004. “Direct Estimation of the Fiber Orientation Density Function from Diffusion-Weighted MRI Data Using Spherical Deconvolution.” NeuroImage 23: 1176–85. https://doi.org/10.1016/j.neuroimage.2004.07.037.

Tuch, David Solomon. 2002. “Diffusion MRI of Complex Tissue Structure.” Massachusetts Institute of Technology.

Wang, R, and Van J Wedeen. 2017. “Trackvis.org.” Martinos Center for Biomedical Imaging, Massachusetts General Hospital.

Winkler, Anderson M, Gerard R Ridgway, Matthew A Webster, Stephen M Smith, and Thomas E Nichols. 2014. “Permutation Inference for the General Linear Model.” NeuroImage 92 (May): 381–97. https://doi.org/http://dx.doi.org/10.1016/j.neuroimage.2014.01.060.

Worsley, K. J., A. C. Evans, S. Marrett, and P. Neelin. 1992. “A Three-Dimensional Statistical Analysis for CBF Activation Studies in Human Brain.” Journal of Cerebral Blood Flow and Metabolism 12: 900–918. https://doi.org/10.1038/jcbfm.1992.127.

Wright, I. C., P. K. McGuire, J.-B. Poline, J. M. Travere, R. M. Murray, C. D. Frith, R. S. J. Frackowiak, and K. J. Friston. 1995. “A Voxel-Based Method for the Statistical Analysis of Gray and White Matter Density Applied to Schizophrenia.” NeuroImage 2 (December): 244–52. https://doi.org/10.1006/nimg.1995.1032.

Yamada, K., K. Sakai, K. Akazawa, S. Yuen, and T. Nishimura. 2009. “MR Tractography: A Review of Its Clinical Applications.” Magnetic Resonance in Medical Sciences 8: 165–74. https://doi.org/10.2463/mrms.8.165.

Yeatman, J. D., R. F. Dougherty, N. J. Myall, B. A. Wandell, and H. M. Feldman. 2012. “Tract Profiles of White Matter Properties: Automating Fiber-Tract Quantification.” PLoS ONE 7. https://doi.org/10.1371/journal.pone.0049790.

Yeh, C.-H., D K Jones, X Liang, M Descoteaux, and A Connelly. 2021. “Mapping Structural Connectivity Using Diffusion MRI: Challenges and Opportunities.” Journal of Magnetic Resonance Imaging 53: 1666–82. https://doi.org/10.1002/jmri.27188.

Yeh, Fang-Cheng, Sandip Panesar, David Fernandes, Antonio Meola, Masanori Yoshino, Juan C. Fernandez-Miranda, Jean M. Vettel, and Timothy Verstynen. 2018. “Population-Averaged Atlas of the Macroscale Human Structural Connectome and Its Network Topology.” NeuroImage 178 (September): 57–68. https://doi.org/10.1016/j.neuroimage.2018.05.027.

Zhang, Fan, Weining Wu, Lipeng Ning, Gloria McAnulty, Deborah Waber, Borjan Gagoski, Kiera Sarill, et al. 2018. “Suprathreshold Fiber Cluster Statistics: Leveraging White Matter Geometry to Enhance Tractography Statistical Analysis.” NeuroImage 171: 341–54. https://doi.org/10.1016/j.neuroimage.2018.01.006.

Zhang, Fan, Ye Wu, Isaiah Norton, Laura Rigolo, Yogesh Rathi, Nikos Makris, and Lauren J. O’Donnell. 2018. “An Anatomically Curated Fiber Clustering White Matter Atlas for Consistent White Matter Tract Parcellation Across the Lifespan.” NeuroImage 179 (October): 429–47. https://doi.org/10.1016/j.neuroimage.2018.06.027.

